# Trehalose and α-glucan mediate distinct abiotic stress responses in *Pseudomonas aeruginosa*

**DOI:** 10.1101/2020.10.23.351924

**Authors:** Stuart D. Woodcock, Karl Syson, Richard H. Little, Despoina Sifouna, James K.M. Brown, Stephen Bornemann, Jacob G. Malone

## Abstract

An important prelude to bacterial infection is the ability of a pathogen to survive independently of the host and to withstand environmental stress. The compatible solute trehalose has previously been connected with diverse abiotic stress tolerances, particularly osmotic shock. In this study, we combine molecular biology and biochemistry to dissect the trehalose metabolic network in the opportunistic human pathogen *Pseudomonas aeruginosa* PA01 and define its role in abiotic stress protection. We show that trehalose metabolism in PA01 is integrated with the biosynthesis of branched α-glucan (glycogen), with mutants in either biosynthetic pathway significantly compromised for survival on abiotic surfaces. While both trehalose and α-glucan are important for abiotic stress tolerance, we show they counter distinct stresses. Trehalose is vital to the PA01 osmotic stress response, with trehalose synthesis mutants displaying severely compromised growth in elevated salt conditions. However, trehalose does not contribute directly to the PA01 desiccation response. Rather, desiccation tolerance is mediated directly by GlgE-derived α-glucan, with deletion of the *glgE* synthase gene compromising PA01 survival in low humidity but having little effect on osmotic sensitivity. Desiccation tolerance is independent of trehalose concentration, marking a clear distinction between the roles of these two molecules in mediating responses to abiotic stress.

## Introduction

*Pseudomonas aeruginosa* is a significant human pathogen and one of the most common causes of nosocomial infections [1]. It is considered a serious medical threat, on a par with methicillin-resistant *Staphylococcus aureus* and extensively drug-resistant *Mycobacterium tuberculosis* [2]. *P*. *aeruginosa* is an opportunistic pathogen that primarily affects immunocompromised individuals, including patients with immunodeficiencies [3], burn or major injury victims [4] and cannula or catheter implant patients [5]. It is also a major pneumonia agent, causing both acute and chronic lung infections. Acute infections typically arise following trauma and can be difficult to eradicate due to a high level of intrinsic antibiotic resistance [6]. Chronic infections often result from a failure to resolve acute infection and involve the persistent, long-term colonisation of the patient [7]. Chronic *P*. *aeruginosa* infections occur in 80 % of adult cystic fibrosis (CF) patients and significantly increase morbidity and mortality compared to uninfected CF patients [7, 8]. The difficulty of treating established *P*. *aeruginosa* infections means that considerable effort is directed towards the prevention of chronic infection in the first place [9].

Throughout its lifecycle *P. aeruginosa* is exposed to diverse environmental challenges, both in association with its hosts and in the wider environment. Outside of an infection, it is subject to frequent changes in temperature, pH, ultra-violet radiation, salinity, humidity and nutrient availability [10, 11]. Similarly, during colonisation of the CF lung *P*. *aeruginosa* is exposed to various abiotic stresses, including nutrient starvation, oxidative stress and desiccation [12]. Similar to other bacteria, *P. aeruginosa* responds to abiotic stresses using a variety of mechanisms, including the formation of biofilms [13], reduction or redirection of metabolic pathways [14], regulation of lipopolysaccharides [15] or stress response proteins [16] and the production of protective solutes [17]. Understanding the mechanisms and pathways *P. aeruginosa* uses to survive abiotic stresses is crucial to reduce and restrict environmental sources of these pathogenic bacteria.

Trehalose is a ubiquitous disaccharide found in virtually all organisms [18] and implicated in diverse stress responses, most notably osmotic stress [19, 20]. In bacteria, trehalose has also been implicated in protection from desiccation stress [21–23], in the development of nitrogen-fixing root nodules [23] and in facilitating bacterial infection in *Mycobacterium tuberculosis* [24, 25] and *Xanthomonas citri* [26]. There are three widespread trehalose metabolic pathways in bacteria [27, 28]. The OtsA/B pathway produces trehalose from glucose 6-phosphate and UDP-glucose via a trehalose-6-phosphate intermediate [29]. A standalone synthase TreS can produce trehalose through the isomerisation of maltose in some metabolic contexts [30]. Finally, the TreY/TreZ pathway produces trehalose by isomerising and hydrolysing α-glucan [28, 31]. Different bacterial species contain varying combinations of all three pathways [32], with most *Pseudomonas* spp. missing OtsA/B, but encoding TreS and TreY/Z.

Bacterial glycogen is a large α-glucan polysaccharide formed from thousands of glucose monomers with α-1,4 glycosidic links and α-1,6 linked branch points [33]. This α-glucan functions as a ubiquitous carbon store, which accumulates during exponential bacterial growth and can be recycled when necessary. It also acts as a virulence factor of *Mycobacterium tuberculosis* [34, 35]. There are two main α-glucan biosynthetic pathways in bacteria. In the classical GlgC-GlgA pathway [36], GlgC generates ADP-glucose from glucose 1-phosphate (G1P) [37, 38]. ADP-glucose is then used by GlgA to extend an α-1,4 linked α-glucan chain [39, 40], with branches being added by GlgB [41, 42]. GlgX and GlgP break down α-glucan by debranching [43] and removing glucose units from the non-reducing end of the glucan [44], respectively. More recently, the alternative GlgE pathway was discovered in *M. tuberculosis* [25] and subsequently found to be encoded in 14% of sequenced bacterial genomes [45]. This pathway employs four enzymes; TreS, Pep2, GlgE and GlgB to produce branched α-glucan from trehalose via maltose and maltose-1-phosphate (M1P) [28, 46], thus making a direct metabolic link between trehalose and α-glucan.

The production and roles of trehalose and α-glucan are currently poorly understood in *Pseudomonas*, with only a handful of studies of trehalose biosynthesis [47, 48] and little knowledge of the role of α-glucan. Recently, deletions of the *treS* and *treY/Z* loci in *P. aeruginosa* PA14 were shown to abolish trehalose production and attenuate model plant infection [48] implicating trehalose as a stress protectant and possible virulence factor. Similarly, trehalose-related gene disruption in *P. syringae* pv. *tomato* resulted in osmotic sensitivity and strains attenuated in plant infection [47]. However, given the potential intrinsic biochemical connections between trehalose and α-glucan biosynthesis, the physiological roles of each molecule in *P. aeruginosa* are currently unclear.

To address this, we fully characterised the trehalose and α-glucan biosynthetic pathways of *P. aeruginosa* to establish how they were configured. We then determined their physiological functions and contribution to abiotic stress tolerance and survival in challenging environments. We demonstrate that contrary to current models, trehalose plays no appreciable role in adapting bacteria to desiccation stress. Rather, this phenotype is mediated in large part by the accumulation of α-glucan. Disruption of either trehalose or branched α-glucan production significantly compromised the ability of *P. aeruginosa* to survive on metal work surfaces, suggesting that abiotic survival in the environment requires the interplay of both stress tolerance phenotypes.

## Results

### Bioinformatic prediction of the trehalose/α-glucan biosynthetic pathway

In *P. aeruginosa* PAO1, the genes associated with trehalose/α-glucan biosynthesis are found in two predicted operons (Figure **1A**) [48, 49]. The predicted functions of the genes are summarised in Table **1**. The first cluster consists of three genes, *glgE*, *treS* and *glgB*, that collectively code for the GlgE α-glucan biosynthetic pathway. The predicted *treS* gene is fused to the predicted maltokinase gene *pep2,* as is most often the case in bacteria containing the GlgE pathway [45], and is henceforth referred to as *treS*/*pep2*. The second cluster encodes classical α-glucan metabolic enzymes [45], consisting of *glgA*, *treZ*, the glucanotransferase gene *malQ* [47], *treY* and *glgX*. The gene for the recycling enzyme GlgP is orphaned and found elsewhere in the genome. In common with other pseudomonads, *P. aeruginosa* does not contain the *otsA/otsB* trehalose synthesis pathway genes [47].

**Figure 1:**
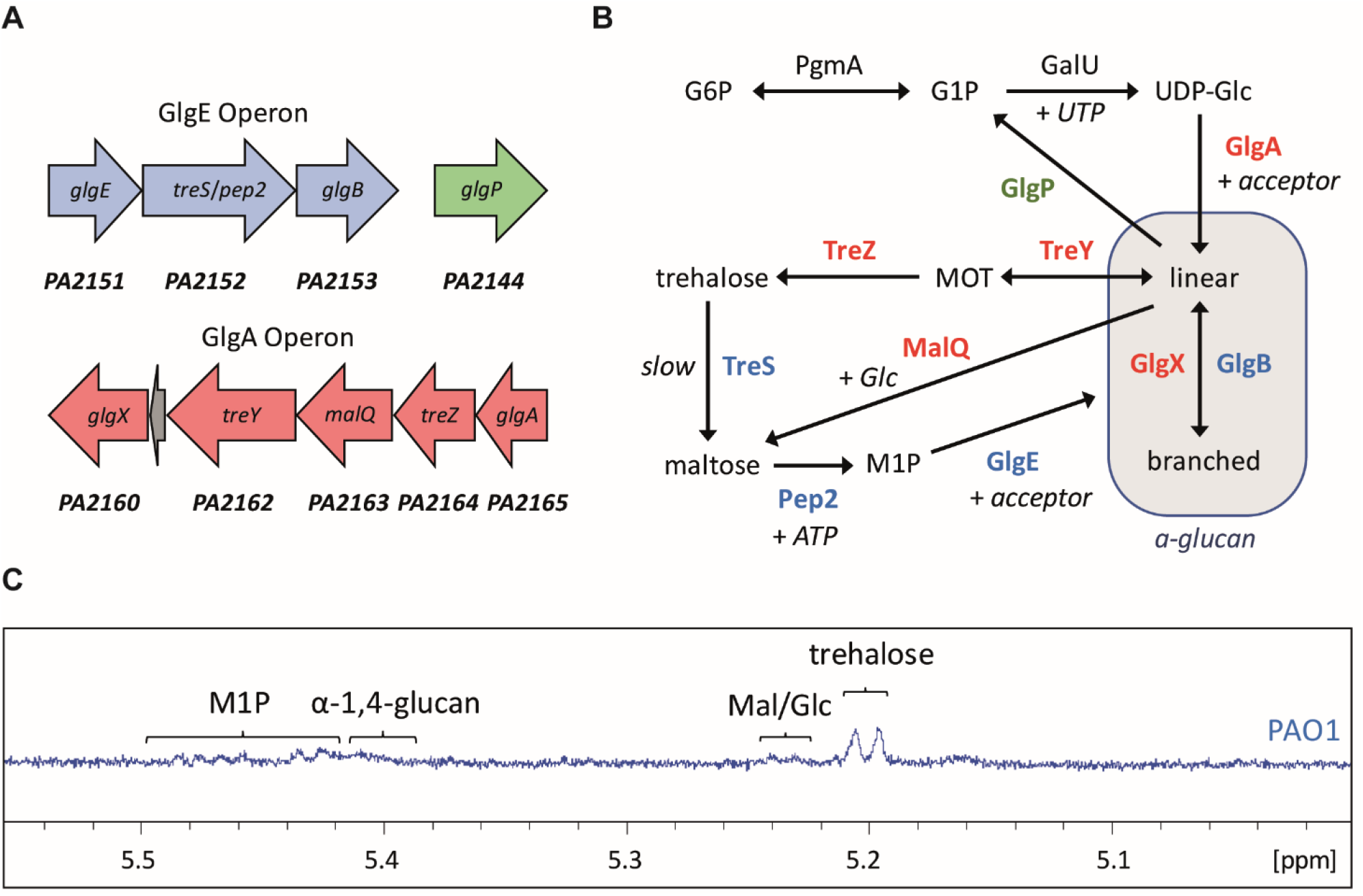
**A) PAO1 trehalose/α-glucan biosynthetic genes** cluster in two predicted operons, while the *glgP* gene is orphaned. Rightward pointing arrows indicate genes on the positive strand, and leftward indicates the negative strand. **B) Model for trehalose and α-glucan biosynthesis in PA01.** The colours of the protein names correspond to the operons shown in 1A. Arrows indicate the primary direction of flux between substrates. Abbreviations are used as follows: G6P – glucose-6-phosphate, G1P – glucose-1-phosphate, UDP-Glc – UDP-glucose, Glc – glucose, MOT – maltooligosyltrehalose, M1P – maltose-1-phosphate. **C) ^1^H-NMR spectrum for PA01.** Peaks corresponding to key metabolites are indicated. Mal/Glc – maltose/glucose.

**Table 1:** Predicted functions of the trehalose/α-glucan metabolic genes in *P. aeruginosa* PA01 that were tested in this study

| Gene name | Gene number | Predicted enzyme function | Reaction catalysed | EC number |
| --- | --- | --- | --- | --- |
| *glgE* | PA2151 | Maltosyltransferase | Extension of α-glucan using M1P as the donor | 2.4.99.16 |
| *treS/*<br>*pep2* | PA2152 | Trehalose-maltose isomerase/maltokinase | Isomerisation between trehalose and maltose/phosphorylation of maltose to M1P using ATP | 5.4.99.16/<br>2.7.1.175 |
| *glgB* | PA2153 | Glycogen branching enzyme | Introduction of α-1,6-linked branches into α-glucan | 2.4.1.18 |
| *glgA* | PA2165 | Glycogen synthase | Extension of α-glucan using UDP-glucose as the donor | 2.4.1.11 |
| *treZ* | PA2164 | Maltooligosyltrehalose hydrolase | Hydrolysis of maltooligosyltrehalose to liberate trehalose | 3.2.1.141 |
| *malQ* | PA2163 | α-Glucanotransferase | Disproportionation of linear maltooligosaccharides with degrees of polymerisation of $\geq 2$ | 2.4.1.25 |
| *treY* | PA2162 | Maltooligosyltrehalose synthase | Conversion of reducing end terminal α-1,4 linkage into an α-1,α-1 linkage, forming maltooligosyltrehalose | 5.4.99.15 |
| *glgX* | PA2160 | Glycogen debranching enzyme | Hydrolysis of α-1,6 linkages of branched α-glucan | 3.2.1.196 |
| *glgP* | PA2144 | Glycogen phosphorylase | Degradation of non-reducing ends of α-glucan yielding glucose 1-phosphate | 2.4.1.1 |

By combining earlier predictions for trehalose biosynthesis in *Pseudomonas* [47, 48] with the known metabolic GlgE pathways in actinomycetes, we produced a model for trehalose/α-glucan biosynthesis in *P. aeruginosa* (Figure **1B**). The clustering of genes within the *glgA* operon suggests it functions to produce trehalose from GlgA-derived linear α-glucan, as previously described [47, 48]. Conversely, the genomic clustering of the four GlgE pathway genes (*glgE*, *treS*/*pep2* and *glgB*) implies a role for this operon in the conversion of trehalose into branched α-glucan, contrary to previous assumptions in pseudomonads. Although PAO1 also contains genes for the phosphomutase PgmA and the UDP-glucose pyrophosphorylase GalU, the ADP-glucose phosphorylase gene *glgC* is not present. This suggests that *P. aeruginosa* uses the activated sugar UDP-glucose as the initial substrate for trehalose/α-glucan biosynthesis [35]. This contrasts with *M. tuberculosis* and most other bacteria that use primarily ADP-glucose [35, 45].

### PAO1 produces α-glucan, trehalose and M1P

To determine the levels of key metabolites in PAO1, the metabolome of the wild-type strain was extracted after growth in M9 medium and analysed using ^1^H-NMR spectroscopy. Wild-type PAO1 accumulated trehalose and M1P to 0.13 ± 0.03 and 0.30 ± 0.03% of the cellular dry weight, respectively. *Pseudomonas* spp. are known to accumulate significantly less α-glucan than other bacteria, with previous studies reporting no detectable α-glucan accumulation for PA14 or *P. syringae Pto* [48, 50]. However, we detected a broad peak corresponding to α-1,4-linkages at 5.41 ppm in the NMR spectrum, consistent with the accumulation of high molecular weight α-glucan in PAO1 (Figure **1C**). Additional evidence for the production of α-glucan is described below. The detection of signals for trehalose, M1P and α-glucan suggests that the GlgE pathway is present and functional in PAO1.

To experimentally test the model in Figure **1B**, in-frame deletions were produced for each of the predicted trehalose/α-glucan biosynthesis genes in PAO1, as well as selected multiple gene deletions. The effect of each gene deletion on the bacterial soluble metabolome was then determined using ^1^H-NMR spectroscopy. The relative accumulations of trehalose and M1P for the tested mutants are summarised in Table **2**, with selected NMR spectra shown in Figure **S1**. The qualitative presence or absence of α-glucan was evident, but quantitation was precluded due to the breadth of the signal. In addition, it was not possible to distinguish glucose and maltose because their α-anomeric signals overlapped, and other distinguishing signals were hidden by signals from other metabolites.

**Table 2:** Accumulation of trehalose and M1P in PAO1 strains. Values are percentages of cellular dry weight ± standard error. – indicates no metabolite detected, * indicates significant differences compared to the wild-type PAO1 as determined by a student’s t-test (p ≤ 0.05, 2-tailed, equal variance). Presence or absence of α-glucan as indicated by + or − respectively, where ++ indicates increased α-glucan production compared with the wild-type strain, ^a^ indicates linear α-glucan only.

| Strain | Trehalose (%) | M1P (%) | Presence of α-glucan |
| --- | --- | --- | --- |
| PAO1 | 0.13 ± 0.03 | 0.30 ± 0.03 | + |
| PAO1 Δ*glgE* | 0.12 ± 0.05 | 2.35 ± 0.43* | ++ |
| PAO1 Δ*treS/pep2* | 0.89 ± 0.25* | - | - |
| PAO1 Δ*glgB* | 0.74 ± 0.08* | 0.78 ± 0.04* | ++<sup>a</sup> |
| PAO1 Δ*glgA* | - | - | - |
| PAO1 Δ*treZ* | - | 0.25 ± 0.06 | ++ |
| PAO1 Δ*malQ* | 0.32 ± 0.04* | 0.22 ± 0.03 | ++ |
| PAO1 Δ*treY* | - | 0.45 ± 0.13 | + |
| PAO1 Δ*glgX* | - | 0.03 ± 0.03* | - |
| PAO1 Δ*glgP* | 0.24 ± 0.01* | 0.29 ± 0.09 | + |
| PAO1 :: *otsA/otsB* | 0.17 ± 0.06 | 0.17 ± 0.02* | + |
| Δ*treS/pep2* :: *otsA/otsB* | 0.91 ± 0.13* | - | - |
| Δ*glgA* :: *otsA/otsB* | 0.06 ± 0.03* | 0.48 ± 0.06* | + |
| Δ*glgA* Δ*glgE* :: *otsA/otsB* | 0.07 ± 0.02 | 0.66 ± 0.11* | - |

### GlgA and GlgP produce and degrade α-glucan chains, respectively

GlgA is annotated as a glycogen synthase [51] and is predicted to be the first committed enzyme in our metabolic model. GlgA is a Glycosyl Transferase 5 family member and is expected to extend linear α-glucan (i.e. maltooligosaccharides) and possibly branched α-glucan using UDP-glucose as the donor substrate. As predicted, deletion of *glgA* led to the complete loss of downstream metabolites in PAO1 (Table **2**). To further test the activity of GlgA, the recombinant protein’s capacity to utilise UDP-glucose was tested *in vitro* using MALDI mass spectrometry. GlgA was able to produce linear α-glucan from UDP-glucose without the addition of an acceptor maltooligosaccharide (Figure **2A**). In addition, the production of α-1,4 linkages was confirmed using NMR spectroscopy with a doublet resonance at 5.32 ppm at the expense of the donor resonances. The most likely priming mechanism involves the slow initial hydrolysis of each NDP-glucose to yield glucose that can subsequently act as the initial acceptor substrate. The highest polymerase activity was with UDP-glucose, as expected given the lack of the *glgC* gene coding for the enzyme responsible for the production of ADP-glucose in *Pseudomonas spp*. GlgA is therefore a UDP-glucose-dependent glycogen synthase whose absence leads to the loss of all trehalose/α-glucan metabolites in PAO1.

**Figure 2:**
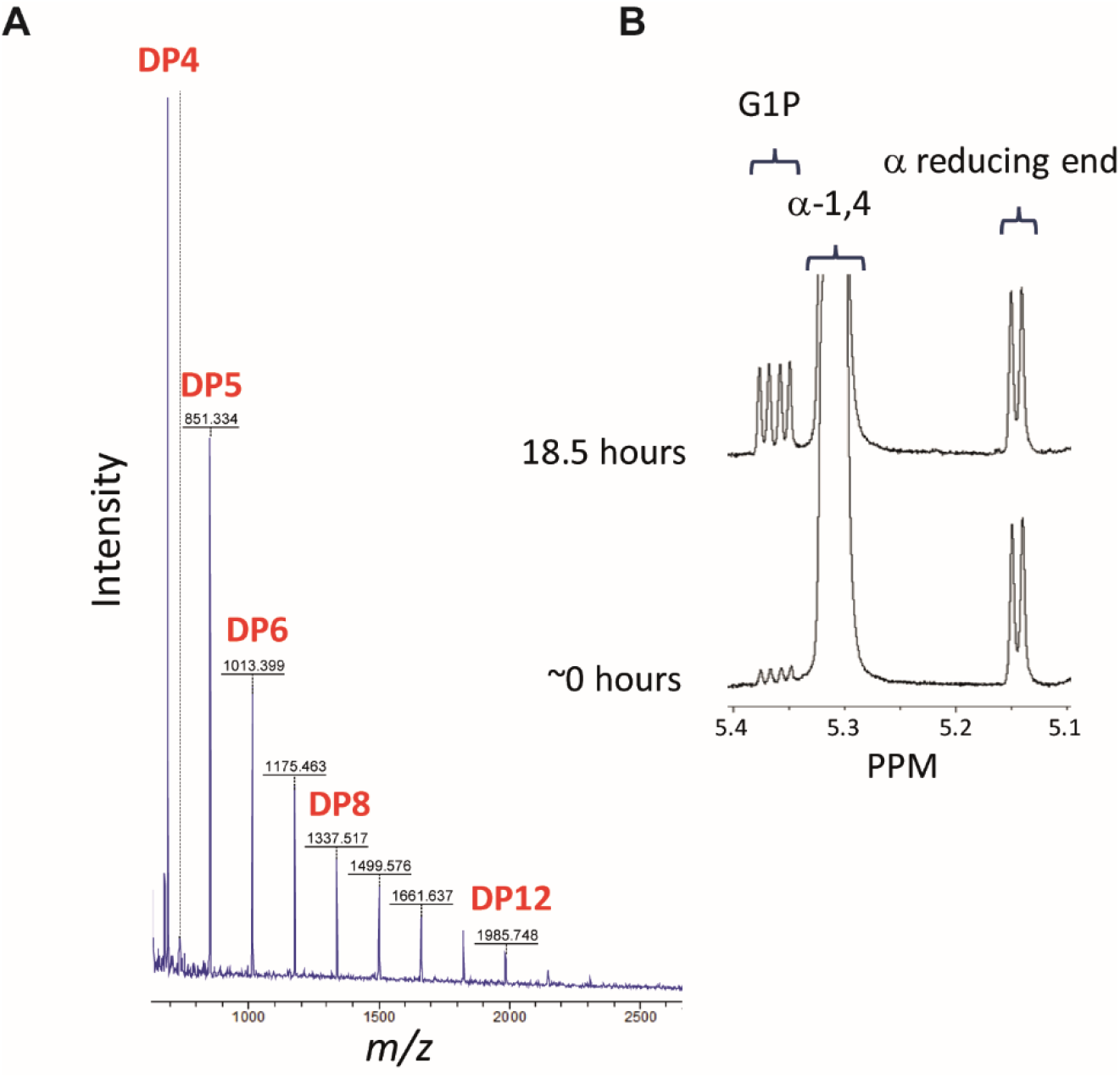
**A) PAO1 GlgA converts UDP-glucose into maltooligosaccharides.** GlgA was incubated with UDP-glucose and the resulting product mixture was analysed using MALDI mass spectrometry. The degree of polymerisation (DP) of the maltooligosaccharide products formed is indicated. **B) PAO1 GlgP is a phosphorylase.** GlgP was incubated with maltohexaose in the presence of inorganic phosphate. The appearance on a ^1^H-NMR spectrum of an anomeric double-doublet resonance at 5.36 ppm is diagnostic of α-glucose-1-phosphate. The shortened maltooligosaccharide co-products maintain the reducing end of the substrate.

Our model suggests that the glycogen phosphorylase GlgP degrades the non-reducing ends of α-glucan into glucose 1-phosphate (G1P). Deletion of *glgP* should therefore result in a higher accumulation of α-glucan and potentially trehalose and M1P. Based on examination of NMR spectra, the level of α-glucan appeared to increase relative to wild-type, although this response could not be quantified. In addition, the level of trehalose increased approximately 2-fold in the Δ*glgP* mutant, although the amount of M1P did not change significantly (Table **2**). To confirm the phosphorylase activity of GlgP, the recombinant protein was incubated with maltohexaose in the presence of inorganic phosphate. NMR spectroscopy showed the expected production of α-glucose-1-phosphate by the appearance of a diagnostic resonance at 5.36 ppm (Figure **2B**). The concentration of reducing ends remained constant, showing that GlgP does not hydrolyse α-1,4 linkages. Thus, GlgP is a glycogen phosphorylase whose absence leads to the increased accumulation of trehalose and α-glucan *in vivo*.

### TreS/Pep2 converts trehalose into maltose-1-phosphate via maltose

Although TreS is commonly annotated as trehalose synthase, net flux through the enzyme will depend on its metabolic context. In the presence of the other GlgE pathway enzymes, flux is predicted to run in the direction of M1P and ultimately branched α-glucan. Such flux is driven by the strongly favourable free energies of the Pep2 maltokinase and branching enzyme reactions, regardless of whether in an *in vitro* [52] or *in vivo* setting [53]. Thus, we predict that the TreS domain of the TreS/Pep2 fusion protein converts trehalose into maltose and the Pep2 domain converts maltose into M1P, driven by the free energy of ATP hydrolysis. Deletion of *treS*/*pep2* should result in an increased level of trehalose and the loss of M1P and other downstream metabolites. As predicted, analysis of the PA01 Δ*treS*/*pep2* metabolome revealed an approximate 7–fold accumulation of trehalose compared to wild-type (Table **2**), alongside the complete loss of M1P and α-glucan. This is fully consistent with TreS/Pep2 converting trehalose into M1P *via* maltose in PAO1.

Recombinant TreS/Pep2 exhibited trehalose synthase and maltokinase activity *in vitro*. Using a coupled assay detecting the co-production of ADP from the Pep2 reaction, activity was detected using either trehalose or maltose as the initial substrate. The pH optimum of the Pep2 maltokinase reaction with trehalose was 8 (Figure **3A**). The rate of this activity increased linearly ~2-fold as a function of temperature between 25 and 50 °C (Figure **3B**). The maltokinase activity conformed to a random order ternary complex mechanism (Figure **3C**), with *k*_cat_ of 43.4 ± 1.5 s^−1^, *K*_m_^ATP^ 0.14 ± 0.01 mM, *K*_m_^Mal^ 0.84 ± 0.08 mM, *K*_i_^ATP^ 0.089 ± 0.021 mM and *K*_i_^Mal^ 0.55 ± 0.13 mM. The activity of the TreS domain was confirmed by observing the equilibration of trehalose-maltose interconversion by NMR spectroscopy when the enzyme was exposed to either trehalose or maltose (5 mM) as the initial substrate. The kinetics of the TreS activity with trehalose was tested by coupling it to the maltokinase assay. This showed that the activity of TreS was potentially rate-limiting with a very high *K*m for trehalose (> 100 mM), giving a *k*_cat_/*K*_m_ of only 0.65 s^−1^ M^−1^ according to the initial slope of an initial rate vs trehalose concentration plot (Figure **3D**).

**Figure 3:**
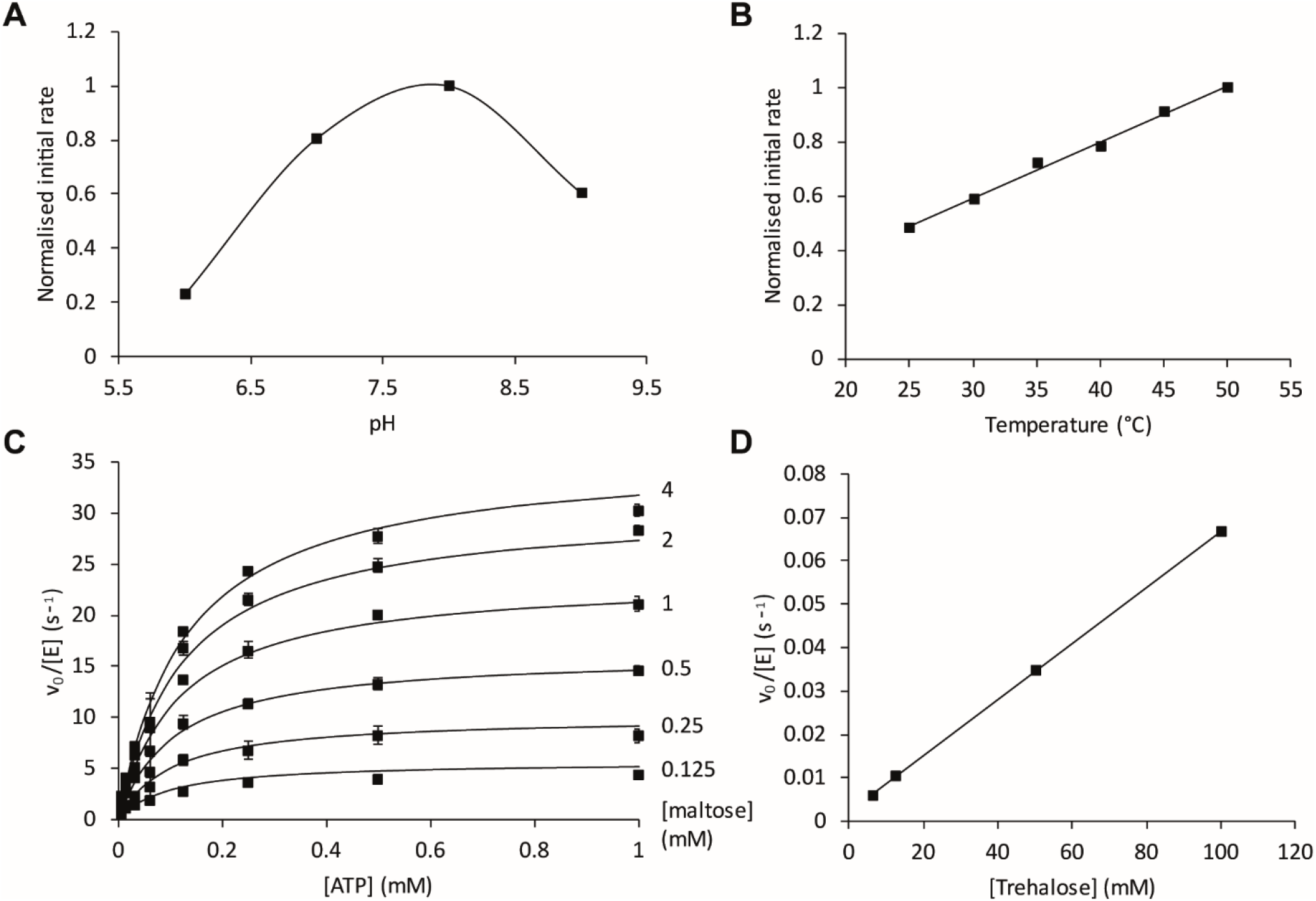
**A-C) Dependence of PAO1 TreS/Pep2 maltokinase activity on pH, temperature, and substrate concentrations.** Maltokinase activity was monitored by detecting the production of ADP using a coupled assay with maltose, phosphoenol pyruvate, pyruvate kinase, lactate dehydrogenase and NADH. **A)** Maltokinase activity of TreS/Pep2 with maltose and ATP was optimal at pH 8. **B)** Maltokinase activity increased ~2-fold linearly with temperature between 25 and 50 °C. Data points in A) and B) represent means of duplicate experiments. **C)** The maltokinase activity of TreS/Pep2 was determined as a function of ATP concentration with a series of fixed maltose concentrations. The smooth lines represent the fit to the data of a random order ternary complex mechanism. Data points show means of triplicate measurements and standard error. **D) The activity of TreS with trehalose** was monitored indirectly by detecting ADP production by Pep2 in a coupled reaction. The *K*m for trehalose was evidently > 100 mM.

### GlgE extends α-glucan with M1P

The maltosyltransferase GlgE is predicted to extend α-glucan using M1P as a substrate (Figure **1**). Deletion of *glgE* should result in the accumulation of M1P and other up-stream metabolites. As predicted, the M1P level increased almost 8-fold to 2.35 ± 0.43% of the cellular dry weight of the PAO1 Δ*glgE* mutant strain (Table **2**). The up-stream metabolites maltose and/or glucose were also detected (not shown), and the (GlgA-derived) α-glucan level also appeared to increase.

Recombinant GlgE extended maltooligosaccharides *in vitro* with the donor M1P, releasing inorganic phosphate. The temperature and pH optima of the enzyme were 30 °C and 7, respectively (Figure **4A** and **4B**). The enzyme preferentially extended linear maltooligosaccharides with a degree of polymerisation (DP) of 5–7 (Figure **4C**). The enzyme exhibited disproportionation activity with maltohexaose, yielding both longer and shorter oligosaccharides by two glucose residues (Figure **4D**) as reported for GlgE enzymes from other bacteria [25, 52]. This side reaction is likely responsible for the substrate inhibition observed with maltohexaose in the polymerase reaction with M1P. Thus, the kinetic constants associated with maltohexaose extension by 1 mM M1P were *k*_cat_^app^ 23 ± 1 s^−1^, *K*_m_^app^ 3.2 ± 0.4 mM and *K*_i_^app^ 171 ± 26 mM (Figure **4E**). With 1 mM maltohexaose, the kinetic constants for M1P were *k*_cat_^app^ 4.3 ± 0.3 s^−1^ and *K*_m_^app^ 0.4 ± 0.1 mM (Figure **4F**). These values are in the same range as those reported for GlgE enzymes from actinomycetes [52].

**Figure 4:**
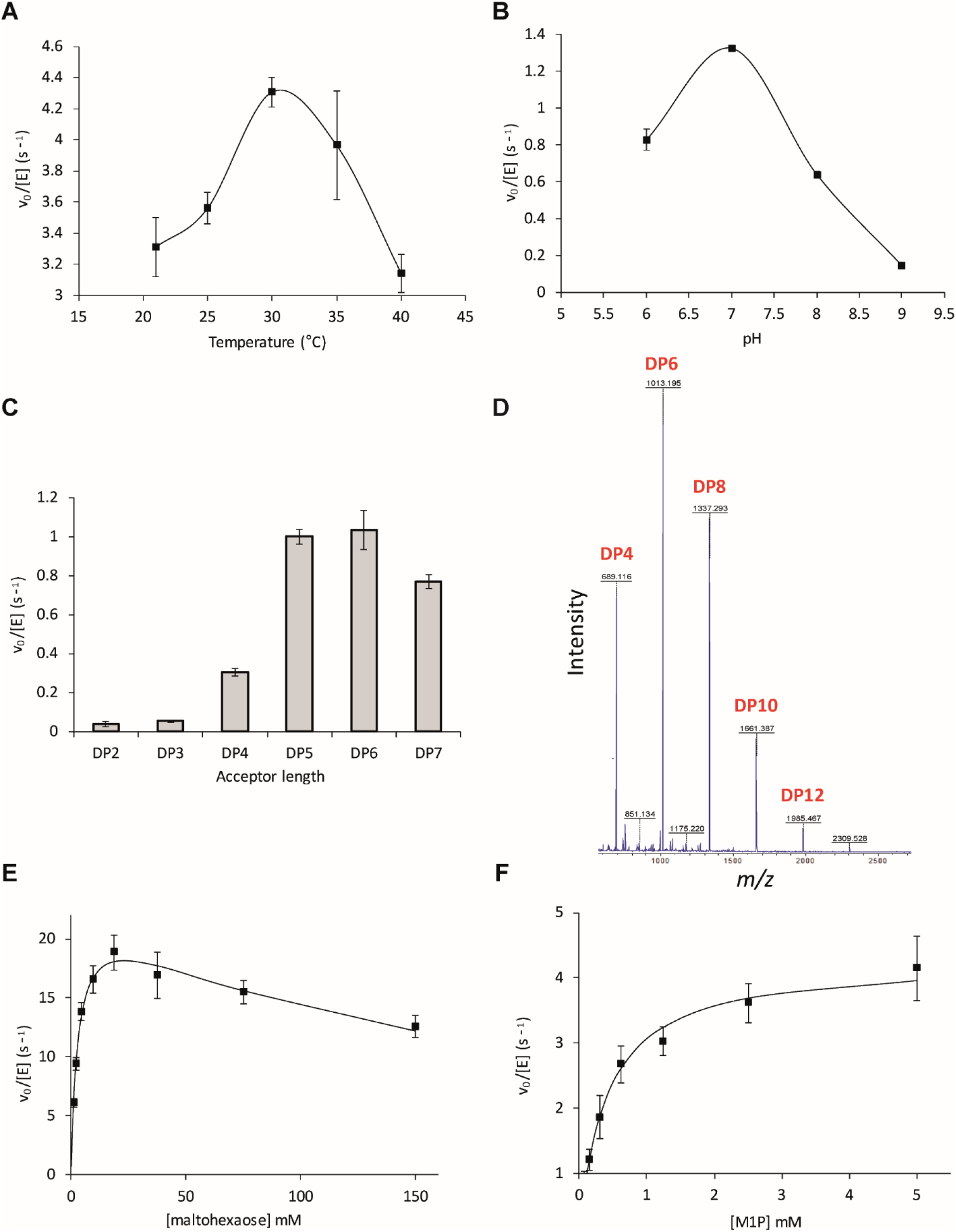
**A-C) Dependence of PAO1 GlgE activity on pH, temperature, and acceptor chain length.** GlgE activity with M1P and an acceptor was monitored by detecting the release of inorganic phosphate in a stopped assay. **A)** The temperature optimum of GlgE activity with maltohexaose was 30 °C. **B)** The pH optimum of GlgE activity with maltohexaose was 7. **C)** GlgE preferentially extends linear maltooligosaccharides with a DP of 5 to 7. All data points represent means of triplicate measurements and standard errors. **D) PAO1 GlgE exhibits disproportionation activity.** GlgE was incubated with maltohexaose and the resulting product mixture was analysed using MALDI mass spectrometry. The DP of the maltooligosaccharide products formed is indicated. Note that GlgE transfers maltose groups between maltooligosaccharides. **E-F) The kinetics of the PAO1 GlgE-catalysed maltohexaose extension by M1P. E)** GlgE activity was determined as a function of maltohexaose concentration using 1 mM M1P. Substrate inhibition was evident at high maltohexaose concentrations. The smooth line represents the fit to the data using the Michaelis-Menten equation modified to allow for substrate inhibition. **F)** GlgE activity was determined as a function of M1P concentration using 1 mM maltohexaose. No substrate inhibition was observed with M1P. The smooth line represents the fit to the data using the Michaelis-Menten equation. All data points represent means of triplicate measurements and standard errors.

### PAO1 TreY and TreZ convert α-glucan into trehalose

TreY is predicted to catalyse the conversion of the reducing-end α-1,4 linkage of α-glucan into an α-1,α-1-linkage, forming maltooligosyltrehalose. Trehalose is in turn predicted to be liberated from maltooligosyltrehalose by the hydrolase TreZ. Deletion of *treY* and *treZ* would be expected to block the production of trehalose leading to an accumulation of GlgA-derived α-glucan. This was indeed observed (Table **2**, Figure **S1**). Deletion of *treZ* alone would be expected to lead to the accumulation of maltooligosyltrehalose. The appearance of a broad NMR signal at 5.2 ppm was consistent with the accumulation of the terminal α-1,1-linkage of this metabolite (Figure **S1**). These data support the prediction that TreY and TreZ together generate trehalose from α-glucan. Unexpectedly, NMR signals corresponding to M1P were present in the PA01 Δ*treY* and Δ*treZ* metabolomes, suggesting that a second, trehalose-independent route exists to form M1P.

### MalQ disproportionates maltooligosaccharides and is capable of synthesising maltose

One possible trehalose-independent route to M1P in the Δ*treY*/*treZ* strain is *via* maltose derived from an activity other than TreS. The most likely candidate in PAO1 is the predicted α-glucanotransferase *malQ*, which is expected to transfer a portion of a maltooligosaccharide with a DP of ≥ 2 to an acceptor with a DP of ≥ 1 . In other words, it would be capable of transferring a glucose group from the non-reducing end of a maltooligosaccharide to a glucose molecule to yield maltose. The recombinant enzyme exhibited disproportionation activity when presented with maltohexaose. The products approached an entropy-driven distribution of products with degrees of polymerisation from DP ≤ 4 to DP ≥ 22 with the shortest products accumulating to the highest levels, according to MALDI mass spectrometry (Figure **5A**). The matrix signals obscured products with a DP of ≤ 3, including maltose with its DP of 2. We therefore incubated MalQ with maltose to establish whether it was capable of interconverting maltose with other maltooligosaccharides. Indeed, it approached an entropy-driven distribution up to a DP of ≥ 10 associated with a proportionately lower average DP, as expected (Figure **5B**). To directly test the hypothesis that α-glucan can be converted to M1P using MalQ and TreS/Pep2, the enzymes were incubated with maltohexaose and ATP and the products were analysed using ^1^H-NMR spectroscopy. M1P was indeed detected after 1 hour (Figure **5C**). In addition, carrying out the reaction in the presence of coupling enzymes to monitor the production of ADP showed the reaction was several orders of magnitude quicker than the background rate of a control in the absence of MalQ. This result shows that MalQ could indeed bypass TreY/TreZ/TreS *in vivo*.

**Figure 5:**
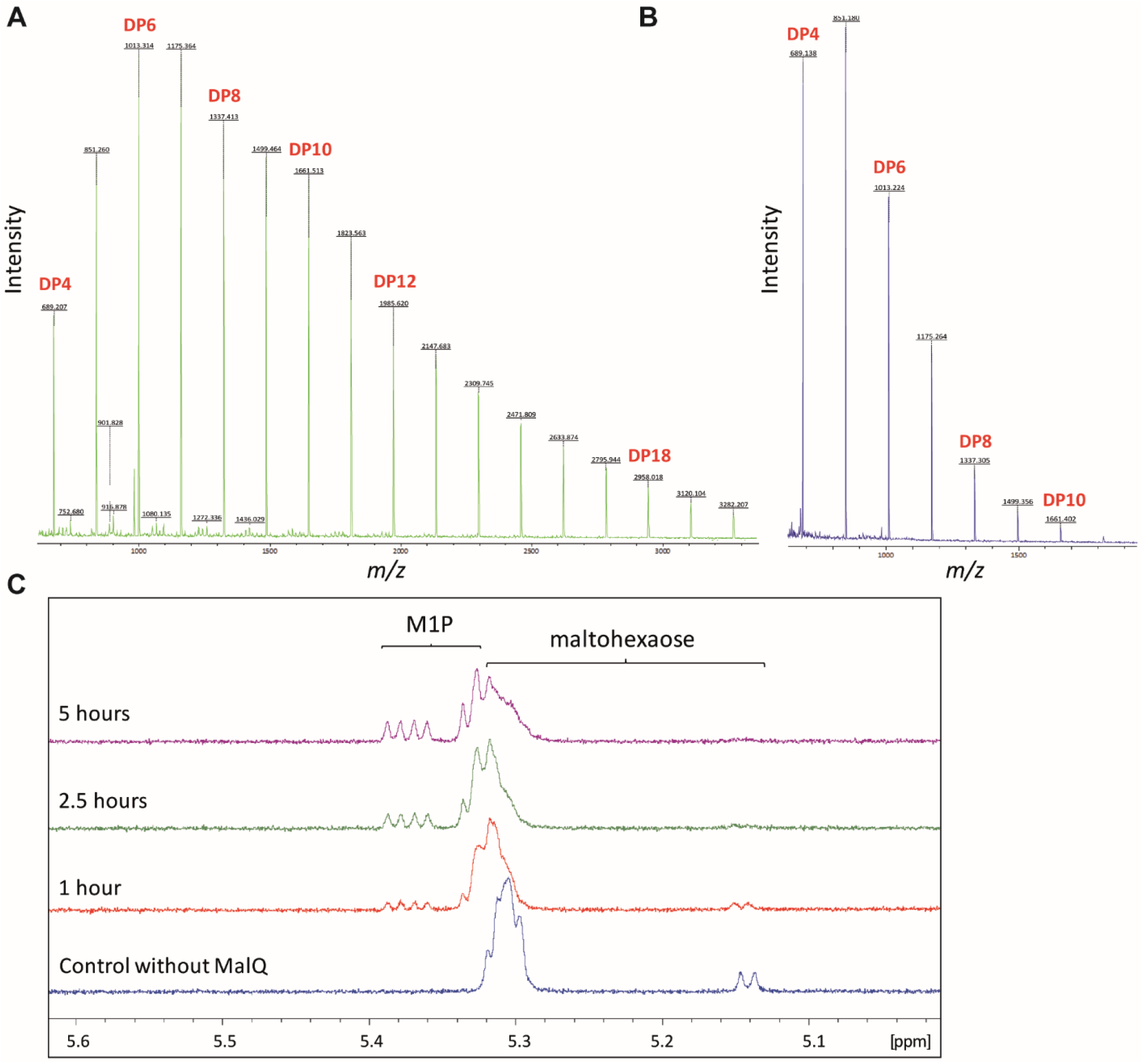
**A) PAO1 MalQ is capable of disproportionating maltohexaose.** MalQ was incubated with maltohexaose and the resulting reaction mixture was analysed using MALDI mass spectrometry. The DP of the maltooligosaccharide products formed is indicated. **B) PAO1 MalQ is capable of disproportionating maltose.** MalQ was incubated with maltose and the resulting reaction mixture was analysed using MALDI mass spectrometry. The DP of the maltooligosaccharide products formed is indicated. **C) Maltohexaose and ATP can be converted to M1P by PAO1 MalQ and TreS/Pep2.** MalQ and TreS/Pep2 were incubated with maltohexaose and ATP and the resulting reaction mixture was analysed using ^1^H-NMR spectroscopy. Diagnostic resonances for M1P appeared concomitantly with the disappearance of those associated with maltohexaose (and other maltooligosaccharide intermediates).

Deletion of the *malQ* gene in PAO1 would therefore be expected to increase the level of GlgA-derived α-glucan leading to increased flux through the TreY/TreZ route to trehalose, and a potentially lower net flux to M1P. The metabolome of this mutant was entirely consistent with these predictions (Table **2**). Thus, MalQ directly contributes to the flux from GlgA-derived α-glucan to GlgE-derived α-glucan, provided glucose is also present. Flux through this route would be enhanced when TreY and/or TreZ are blocked leading to an increase in the production of the MalQ α-glucan substrate derived from GlgA.

### GlgB and GlgX introduce and hydrolyse α-1,6-linked branches in α-glucan particles, respectively

The branching enzyme GlgB is predicted to introduce branches into α-glucan by transferring a portion of a maltooligosaccharide to the 6-OH position of the same or a different α-glucan. Deletion of *glgB* should result in the absence of branched α-glucan and an accumulation of longer linear α-glucan chains and upstream metabolites. Indeed, PAO1 Δ*glgB* showed an increased level of trehalose, maltose/glucose, M1P (Table **2**) and, (presumably linear) α-glucan (Figure **S1**). To assess the length of linear α-glucan chains in PAO1 strains, we exposed colonies to iodine vapour [54]. Short linear chains do not stain, and branched polymers give a pale red/brown colour with iodine. PAO1 gave slight brown staining when exposed to iodine consistent with the production of branched α-glucan. By contrast, the *glgA* mutant gave essentially no staining consistent with a lack of α-glucan. Importantly, the Δ*glgB* strain stained a much darker colour than the wild-type strain, consistent with the presence of significant amounts of long linear α-glucan (Figure **6A**). Similarly, the addition of Lugol’s solution to the metabolite extract from the Δ*glgB* strain gave a dark purple colour associated with linear α-glucans with a DP of between 33 to 38 that are barely soluble (Figure **6B**) [54]. Not only does this confirm the branching activity of GlgB, but also the production of α-glucan *in vivo*.

**Figure 6:**
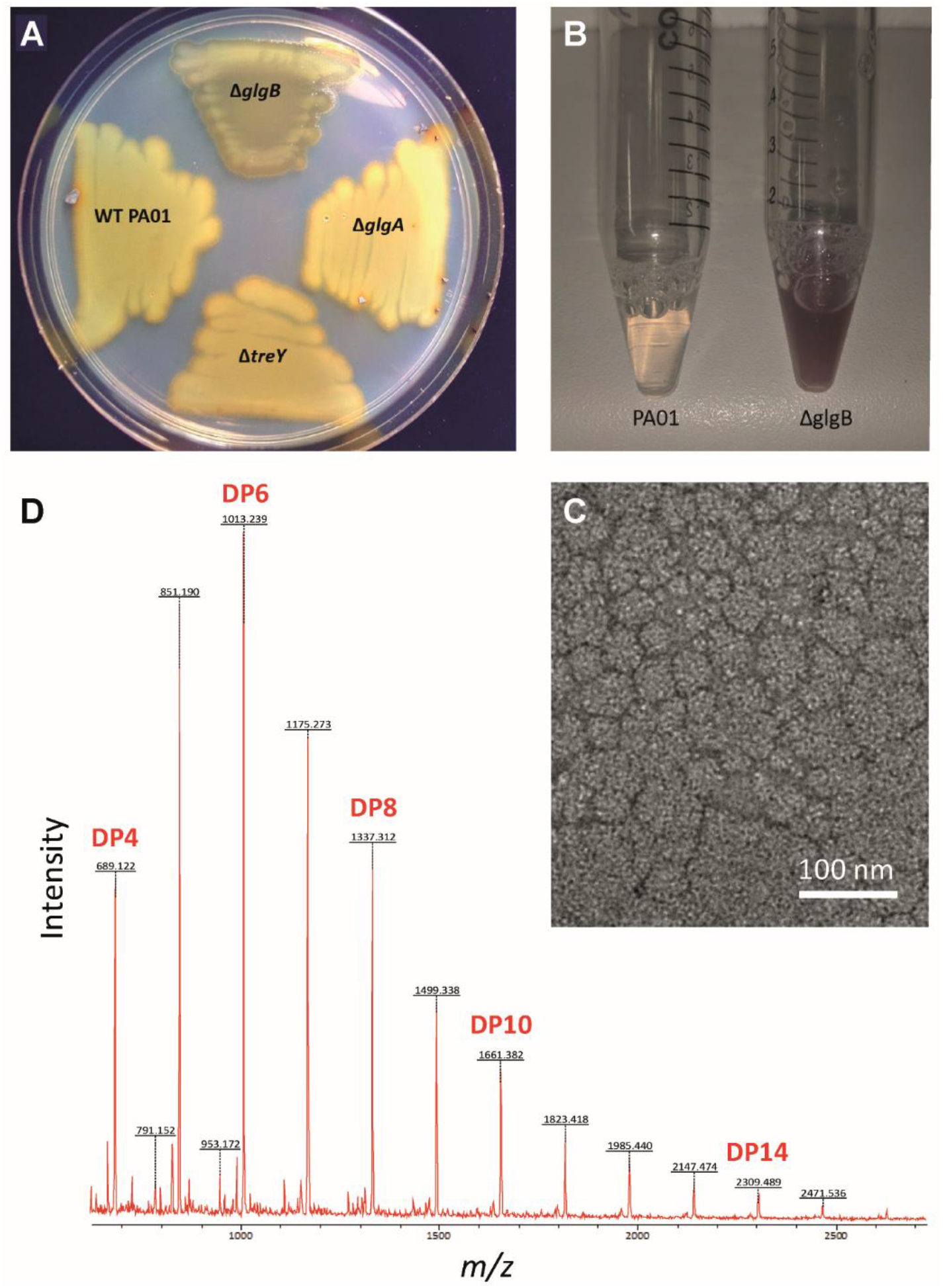
**A) PA01 strains exposed to iodine vapour.** Δ*glgB* stains darker than the other tested strains due to the presence of insoluble linear α-glucan. **B) Lugol’s solution** stains the Δ*glgB* sample dark purple **C) PAO1 GlgB and GlgE generate particles from maltohexaose and M1P.** GlgB and GlgE were incubated with maltohexaose and M1P and the resulting reaction mixture was analysed using transmission electron microscopy. Particles with diameters of tens of nm are evident. **D) Linear chain lengths of synthetic α-glucan.** The α-glucan particles generated by PAO1 GlgB and GlgE were exhaustively unbranched using isoamylase and analysed by MALDI mass spectrometry. The average DP of the chains was 7.6 ± 2.5 with a mode of 6.

Recombinant PAO1 GlgB activity was investigated *in vitro* by incubating it with recombinant PAO1 GlgE, maltohexaose and M1P. Transmission electron microscopy of the resulting product mixture showed the presence of particles with diameters ranging from 20 to over 100 nm, with most around 40 nm (Figure **6C**). These sizes are similar to those observed for particles of α-glucan isolated from other bacteria or generated *in vitro* with GlgE and GlgB enzymes from other bacteria [55].When the particles were hydrolysed using an isoamylase that cleaves all of the α-1,6 linkages, MALDI-MS revealed that the polymer was composed of linear chains with a mean DP of 7.6 ± 2.5 with a mode of 6 (Figure **6D**). The chain length of α-glucan is largely dictated by the branching enzyme GlgB in such systems [55]. Importantly, these experiments show that the combination of GlgE and GlgB from PAO1 is capable of generating branched α-glucan particles.

The glycogen debranching enzyme GlgX is predicted to hydrolyse the α-1,6 links in branched α-glucan. Blocking this enzyme would severely restrict the recycling of branched α-glucan because the other enzymes that degrade the polymer can only operate on the α-1,4 linkages of external chains. For example, although GlgP can in principle operate on all of the non-reducing ends of the arboreal α-glucan polymer, it can only do so until it gets to within a few glucose residues of the point where each external branch is attached to the rest of the polymer via an α-1,6 linkage. TreY and TreZ are even more restricted because they operate on reducing ends, of which an α-glucan polymer has only one. Furthermore, the reducing end can be sterically hindered by the rest of a mature polymer particle. Given the thermodynamics of the α-glucan pathways, a PAO1 Δ*glgX* mutant would be expected to accumulate branched α-glucan (i.e. a metabolic sink) at the expense of upstream metabolites. As predicted, the Δ*glgX* mutant showed a significant reduction in the levels of trehalose and M1P (Table **2**). Unexpectedly, we saw no obvious signal for α-glucan in the Δ*glgX* mutant.

To examine the activity of recombinant PAO1 GlgX, it was incubated with rabbit liver glycogen composed of linear chains with a DP of 10.8 ± 4.6 [55]. MALDI-MS showed that the enzyme preferentially liberated a maltooligosaccharide with a DP of 4 and significantly less material with a DP of 5 and above (Figure **7**), supporting its role as a debranching enzyme with a short chain-length specificity.

**Figure 7:**
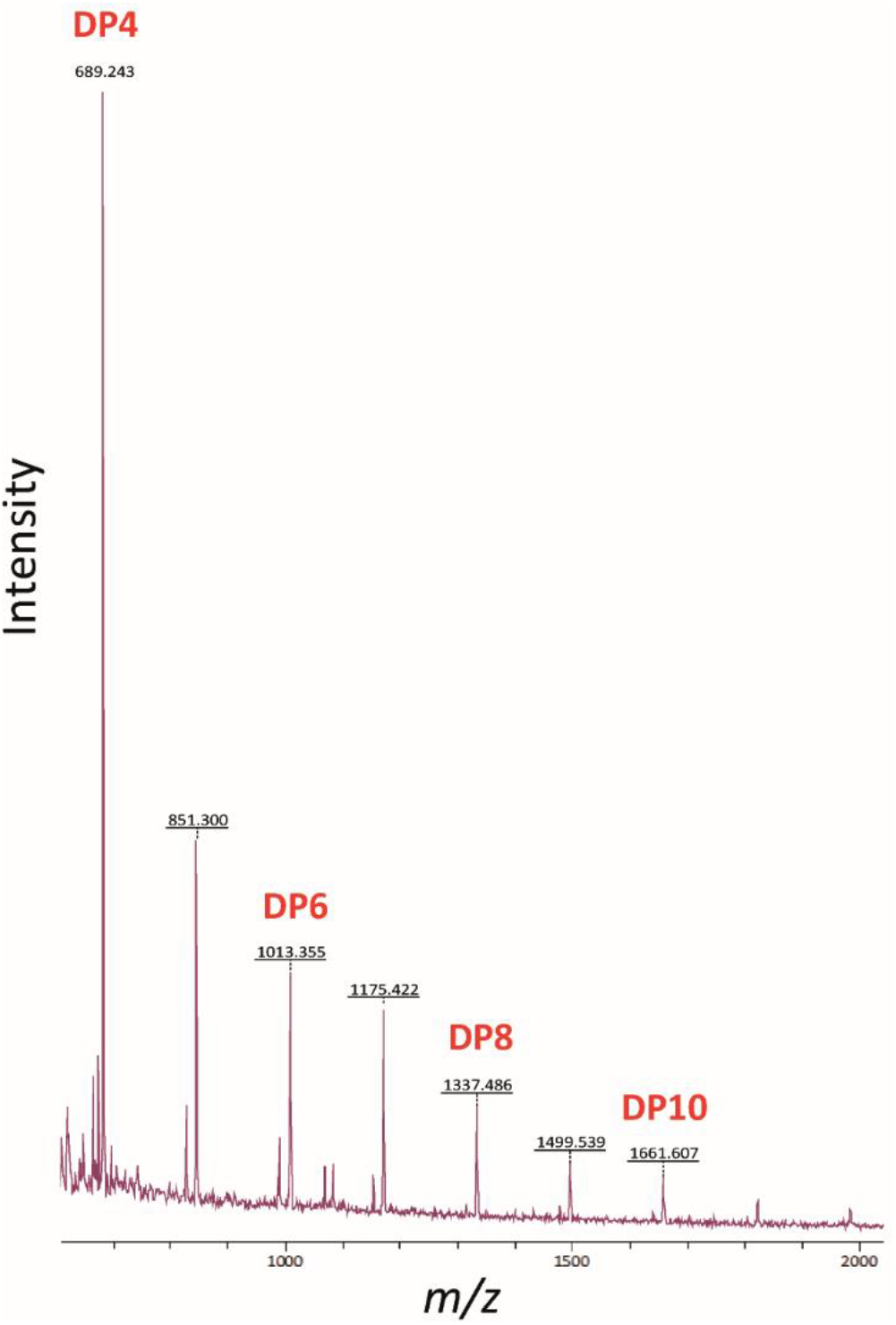
PAO1 GlgX is a debranching enzyme with a short chain length specificity. GlgX was incubated with rabbit liver glycogen that is composed of linear chains with an average DP of 10.8 ± 4.6. The resulting reaction mixture was analysed using MALDI mass spectrometry. The DP of the liberated linear maltooligosaccharide products is shown, with maltotetraose being the major product detected.

### Distinguishing between linear α-glucan pools

At this stage it was unclear whether the linear α-glucan pools produced by GlgA or GlgE are structurally/ spatially distinct or if they are interchangeable. To try to distinguish between these two possibilities, we produced a GlgE pathway deletion mutant (i.e. missing the entire cluster comprising *glgE*, *treS/pep2* and *glgB* - Δ*PA2151-53*). Based on iodine staining (Figure **S2**), this strain accumulated no more α-glucan than wild-type PA01 (Figure **S2A, S2C**). To test whether MalQ modified the GlgA-derived α-glucan, the *malQ* gene was deleted in the GlgE pathway mutant strain background. However, no significant change in iodine staining was apparent in the resulting strain (Figure **S2E**). To test whether the GlgA-derived α-glucan pool was rapidly converted to trehalose, *treY* was deleted in the GlgE pathway mutant background (PA01 Δ*4*). The significant blue staining of this strain with iodine (Figure **S2D**) showed that it accumulated long insoluble α-glucan chains with a DP of over 40 [54], longer than those detected in the *glgB* mutant. Thus, GlgA is capable of generating long linear α-glucan chains *in vivo* (with DPs that might be modified by MalQ). However, such GlgA-derived α-glucan does not accumulate in the cell unless its conversion to trehalose by TreY and TreZ is blocked. Overexpression of *glgB in trans* abolished the blue staining of the PA01 Δ*4* mutant when exposed to iodine (Figure **S2F**), indicating that GlgA-derived α-glucan could be branched, and thus solubilised by GlgB, similarly to wild-type α-glucan.

### Disruption of trehalose biosynthesis results in osmotic sensitivity in PAO1

Bacterial water stress manifests as at least two distinct phenomena: osmotic and desiccation stress [17]. Desiccation stress is distinct from osmotic stress in that it is independent of solute concentration: bacteria are exposed to air and water is lost to the atmosphere and cannot be recovered [56]. There is substantial overlap in the bacterial responses to these two stressors, although efforts have been made to distinguish individual responses from each other [57]. Trehalose has previously been associated with both osmotic and desiccation stress tolerance in *Pseudomonas* spp. [47, 48]. However, the newly revealed connection between trehalose and α-glucan metabolism begs the question as to whether α-glucan plays a direct role in mediating these phenotypes.

To investigate the association of trehalose and/or α-glucan with osmotolerance in PAO1, the growth of selected *glg* mutants were compared in the presence of elevated concentrations of salt (Figure **8**). PA01 grown in M9 minimal medium containing 0.85 M NaCl exhibited a severely delayed lag-phase but higher maximal density compared to growth without added NaCl (Figure **8A**). PAO1 Δ*glgA* (trehalose^−^/α-glucan^−^) showed an even longer lag phase in the presence of salt than wild-type PA01, consistent with trehalose and/or α-glucan conferring osmotolerance (Figure **8B**). The Δ*treS*/*pep2* (trehalose^++^, GlgE α-glucan^−^), Δ*glgE* (GlgE α-glucan^−^, Figure **8A**), and Δ*malQ* (trehalose^+^, Figure **8B**) strains grew more similarly to the wild-type strain, implying trehalose rather than GlgE-derived α-glucan offered protection. Similarly, PAO1 Δ*treY* and Δ*treZ* (trehalose^−^, Figure **8B**) showed significantly attenuated growth in elevated salt. Since the accumulated α-glucan in several of these strains would be expected to derive more from GlgA than GlgE, this result implies that the GlgA-derived α-glucan does not contribute to osmotolerance. The significant attenuation of the Δ*glgX* (trehalose^−^, Figure **8B**) mutants again supports the role of trehalose in osmo-protection. Interestingly, the PA01 Δ*glgB* mutant (trehalose^+^, Figure **8A**) was also strongly attenuated in high-salt growth, despite this strain accumulating more trehalose than the wild-type strain (Table **2**). This suggests that linear, insoluble α-glucan not only does not contribute to osmotolerance but may actually impede an effective osmostress response, perhaps due to reduced potential for metabolism to trehalose.

**Figure 8:**
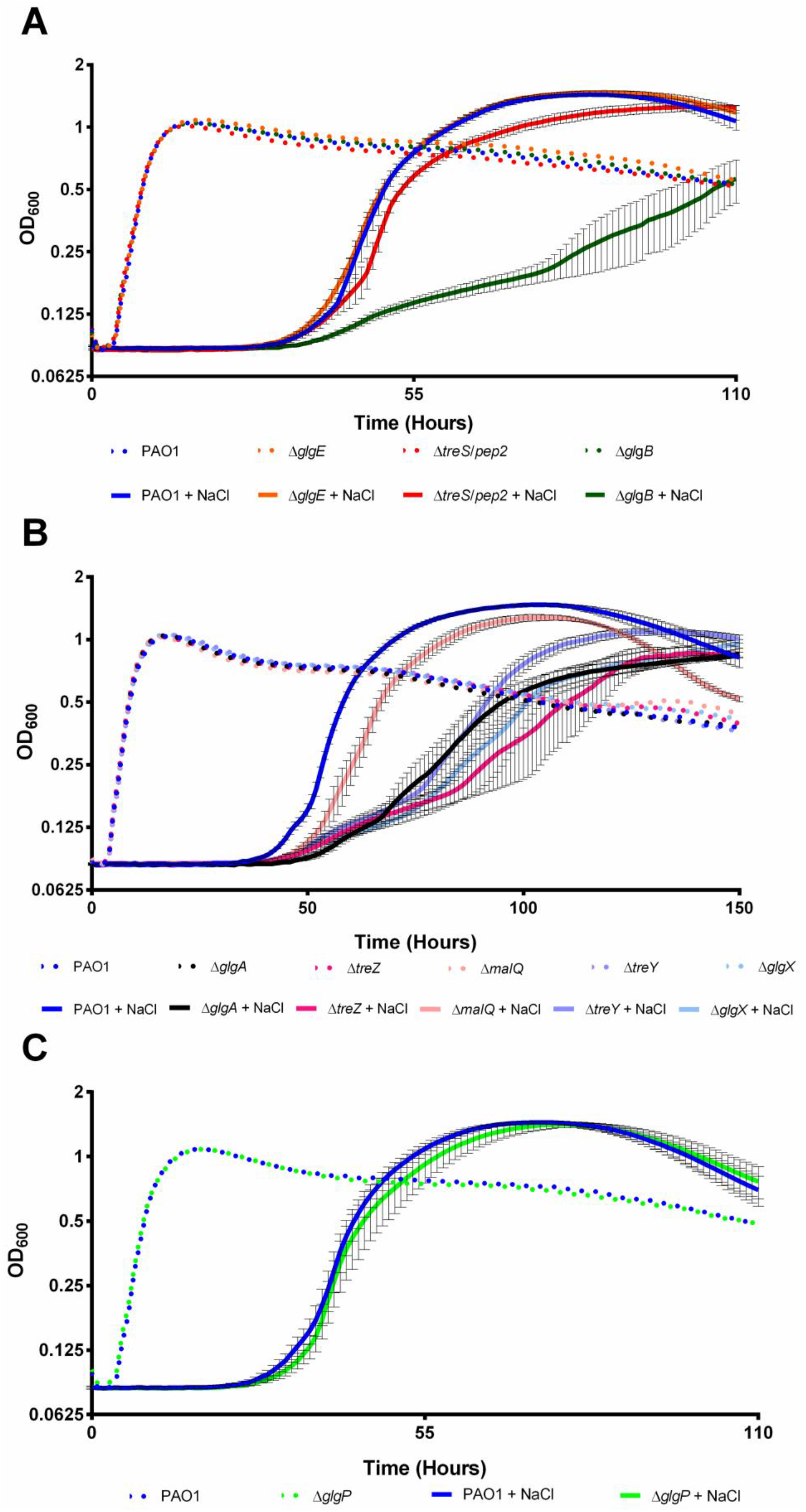
Growth curves of PA01 mutant strains subject to osmotic stress conditions. OD_600_ is plotted against time for strains grown in the presence (+ NaCl, solid lines) and absence (dotted lines) of 0.85 M NaCl. Data are shown +/− standard error of 3 technical replicates. At least two independent biological replicates were conducted per strain and a representative curve is shown in each case. **A)** Wild-type (WT) PA01, Δ*glgE,* Δ*treS*/*pep2* & Δ*glgB.* **B)** WT PA01, Δ*glgA,* Δ*treZ,* Δ*malQ,* Δ*treY* & Δ*glgX* **C)** WT PA01 & Δ*glgP*.

### Loss of GlgE-derived α-glucan, but not trehalose, results in PA01 desiccation sensitivity

Next, we assessed the ability of PA01 mutants to survive exposure to prolonged periods of low relative humidity (RH). Colony forming units (CFU) recovered after exposure of bacterial samples to 100% and 75% RH were counted, with the resulting data analysed using linear mixed modelling and represented as predicted means of log_10_(CFU/ml). The sensitivity of a strain to lower humidity was calculated as the difference between its predicted mean log_10_(CFU/ml) at 100% and at 75% RH. A greater response to lower RH of a mutant compared to the wild-type translates to a more desiccation-sensitive strain. Incubation of all PA01 strains at 100% RH resulted in the recovery of predicted means of 7.5 – 7.9 log_10_(CFU/ml). Incubation at 75% RH led to an approximately 10-fold reduction in the recovery of wild-type PAO1 to a predicted mean of log_10_(CFU/ml) = 6.5 (Figure **9A**), translating to a desiccation response of ~1.1 log_10_(CFU/ml).

**Figure 9:**
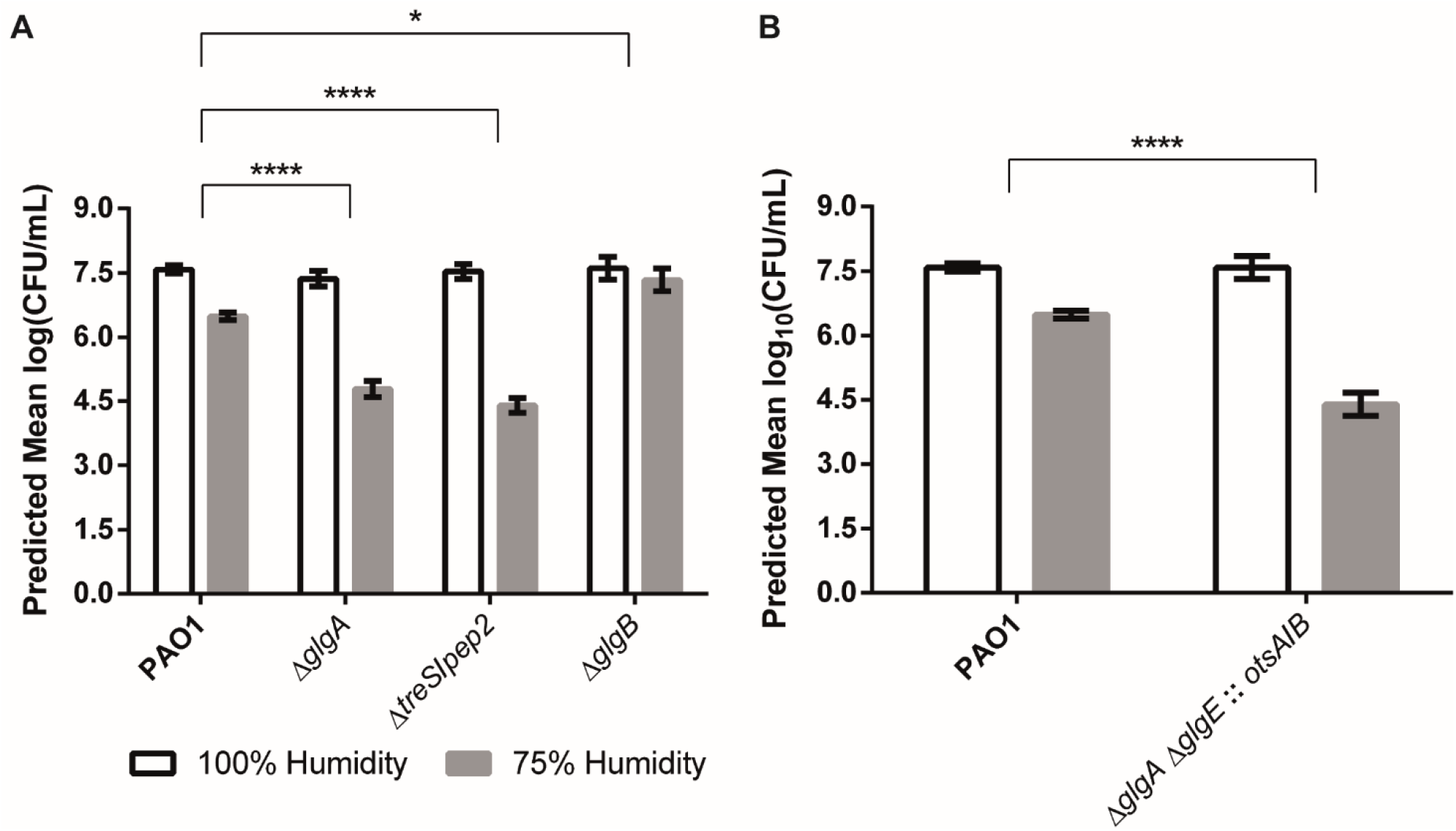
Survival of PA01 mutant strains subject to desiccation stress conditions. Graphs show predicted mean log(CFU/ml) for strains grown in 100% (white bars) or 75% (grey bars) relative humidity. The response to desiccation is the difference log(CFU/ml) at 100% and 75% RH. P-values for differences between strains’ responses were calculated by linear mixed modelling and are shown with asterisks (*: 0.05>p>0.01, ****: p<0.0001). Responses were tested in at least two independent biological replicates on different dates. **A)** Strains lacking GlgE-derived α-glucan (Δ*glgA,* Δ*treS/pep2*) are desiccation sensitive. Glucan branching (by GlgB) is not required for desiccation tolerance. **B)** Recovery of trehalose synthesis by *otsA/B* expression cannot recover desiccation tolerance in a Δ*glgA/E* (α-glucan minus) mutant.

It has previously been reported that trehalose contributes to desiccation tolerance in *P. aeruginosa* [22]. A Δ*glgA* mutant lacking all trehalose (and α-glucan) would therefore be expected to be more sensitive to desiccation than the wild-type strain. Incubation of PAO1 Δ*glgA* at 75% RH resulted in enhanced cell death with a reduction in cell recovery to 4.8 log_10_(CFU/ml)) and a desiccation response of ~2.6 log_10_(CFU/ml). Surprisingly, the trehalose-producing, GlgE-derived α-glucan-null mutant PAO1 Δ*treS*/*pep2* showed a similar sensitivity to the Δ*glgA* mutant, with a desiccation response of ~3.1 log_10_(CFU/ml) (Figure **9A**). This was despite this strain having a trehalose level nearly 7-fold higher than the wild-type strain (Table **2**). This suggested that GlgE-derived α-glucan protects against desiccation stress. Interestingly, the PAO1 Δ*glgB* mutant also showed significantly (p <0.05) enhanced desiccation tolerance, with a desiccation response of ~0.3 log_10_(CFU/ml), implying the GlgE-derived α-glucan did not need to be branched to afford protection (Figure **9A**).

Overexpression of the *otsA* and *otsB* trehalose biosynthetic genes from *M*. *koreensis* 3J1 has previously been shown to increase trehalose production and confer increased desiccation resistance in *P*. *putida* [22, 58]. These genes were introduced into a strain lacking both α-glucan polymerases (GlgA and GlgE) so that it was able to produce trehalose and, given the presence of TreS/Pep2, also maltose and M1P. This strain showed desiccation sensitivity comparable to the strain lacking TreS/Pep2, with a desiccation response of 3.2 log_10_(CFU/mL) (*p* ≤ 0.001) (Figure **9B**). This was despite the trehalose level (0.07 ± 0.02%, Table **2**) being similar to that of wild-type PAO1. A significant ~2-fold increase in the M1P level to 0.66 ± 0.11% (*p* ≤ 0.05) of cellular dry weight compared with the wild-type strain suggests that M1P does not contribute to desiccation stress tolerance in PAO1. Together, these results indicate that GlgE-derived α-glucan is a key mediator of *P. aeruginosa* desiccation tolerance, whether branched or not, and that trehalose does not directly offer protection.

### Trehalose correlates with osmotolerance, while α-glucan correlates with desiccation tolerance

Our data show that the nearly 7-fold higher trehalose level in PAO1 Δ*treS*/*pep2* did not protect this strain against desiccation stress. This unexpected observation led us to explore the impact of increasing the level of trehalose in other strains on osmotolerance and desiccation tolerance. Consequently, *M*. *koreensis otsA* and *otsB* were constitutively expressed in PAO1 wild-type, Δ*glgA* and Δ*treS*/*pep2* strains and their effect on bacterial behaviour and metabolism was examined.

Expression of *otsA*/*otsB* in PAO1 resulted in increased trehalose accumulation and a loss of M1P compared to wild-type PAO1 (Figure **10A**). OtsA/OtsB and GlgA compete for the shared substrate UDP-glucose, so the GlgA-derived α-glucan level would be expected to decrease in this strain. In Δ*glgA*, OtsA/OtsB restored trehalose, M1P and α-glucan biosynthesis, which would be exclusively derived from GlgE. Expression of *otsA*/*otsB* in Δ*treS*/*pep2* resulted in a small increase in trehalose compared to the parental Δ*treS*/*pep2* mutant, while M1P and α-glucan could not be detected, as observed with Δ*treS*/*pep2* (Table **2**). Expression of *otsA*/*otsB* did not affect the growth of any of these strains when no stress was applied. However, they were all strongly enhanced in their ability to tolerate osmotic stress compared to their respective parental strains lacking OtsA/OtsB (Figure **10B**). This further supports the hypothesis that trehalose, rather than α-glucan, protects against osmotic stress in *Pseudomonas*.

**Figure 10:**
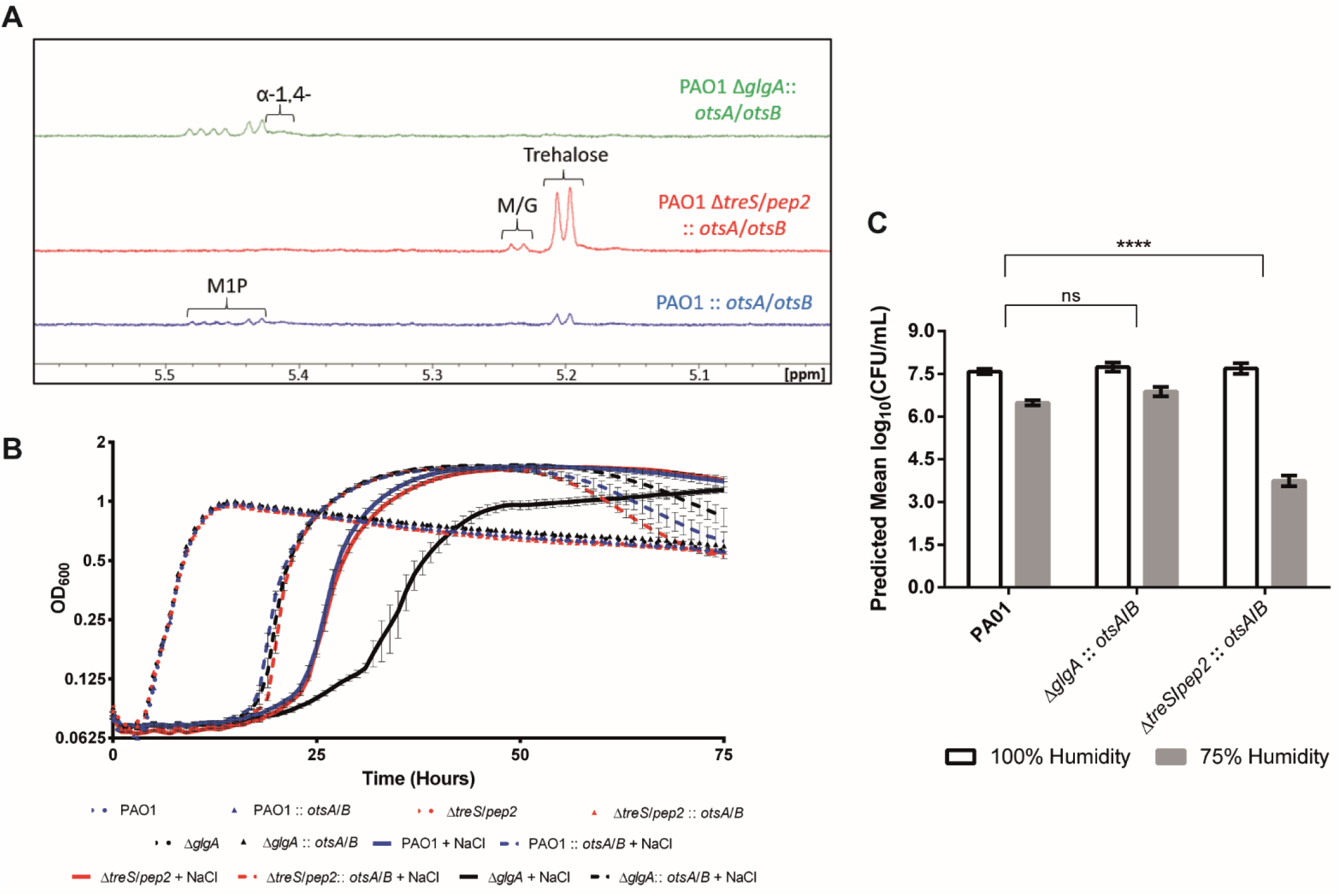
**A) ^1^H-NMR spectra for PA01 strains expressing** *otsA/B*. Peaks corresponding to key metabolites are indicated. M/G – maltose/glucose, α-1,4-– α-glucan. B) **Growth curves of PA01 strains expressing** *otsA/B* **subject to osmotic stress conditions.** OD_600_ is plotted against time for strains grown in the presence (+ NaCl, solid lines) and absence (dotted lines) of 0.85 M NaCl. Data are shown +/− standard error of 3 technical replicates. At least two independent biological replicates were conducted per strain and a representative curve is shown in each case. **C) Survival of PA01 strains expressing** *otsA/B* **subjected to desiccation stress conditions.** Graphs show predicted mean log(CFU/ml) for strains grown in 100% (white bars) or 75% (grey bars) relative humidity. The response to desiccation is the difference log(CFU/ml) at 100% and 75% RH. P-values for differences between strains’ responses were calculated by linear mixed modelling and are shown with asterisks (ns: p>0.05 [not significant], ****: p<0.0001). Responses were tested in at least two independent biological replicates on different dates.

The expression of *otsA*/*otsB* rescued the desiccation sensitivity exhibited by PAO1 Δ*glgA* (Figure **10C**), resulting in a desiccation response of 0.9 log_10_(CFU/mL). This showed that either trehalose or GlgE-derived α-glucan confer tolerance to desiccation. In contrast, PAO1 Δ*treS*/*pep2*::*otsA*/*otsB* exhibited a desiccation response of approximately 4 log_10_(CFU/mL) (*p* ≤ 0.001) translating to significant desiccation sensitivity compared to wild-type, despite accumulating trehalose to above wild-type levels (Figure **10A**). Together our results strongly suggest that trehalose is not directly responsible for desiccation resistance in *Pseudomonas* spp. but rather exerts its desiccation tolerance effect indirectly through the biosynthesis of GlgE-derived α-glucan.

### Trehalose and α-glucan contribute to *P. aeruginosa* survival on abiotic surfaces

Finally, we assessed the contribution of trehalose and α-glucan biosynthesis to *P. aeruginosa* survival in the wider environment. We sought to mimic the conditions these bacteria might encounter upon contaminating a kitchen countertop or other work surface. Therefore, 10 μl droplets of PA01 strains were allowed to dry onto a sterile, brushed-steel worksurface, then recovered and quantified after 24h incubation at room temperature. Alongside wild-type PAO1 we examined the relative survival of Δ*glgA* (trehalose^−^, α-glucan^−^), Δ*glgB* (trehalose^+^, no branched glucan), Δ*treY* (trehalose^−^) and Δ*treS*/*pep2* (trehalose^+^, little/no α-glucan). These four strains were chosen to represent a range of different trehalose and α-glucan abundances and desiccation/osmostress tolerance states. Surprisingly, all four mutants were significantly compromised for surface survival compared with wild-type PAO1 (Figure **11**), supporting roles for both trehalose and α-glucan biosynthesis in mediating *P. aeruginosa* survival on abiotic surfaces.

**Figure 11:**
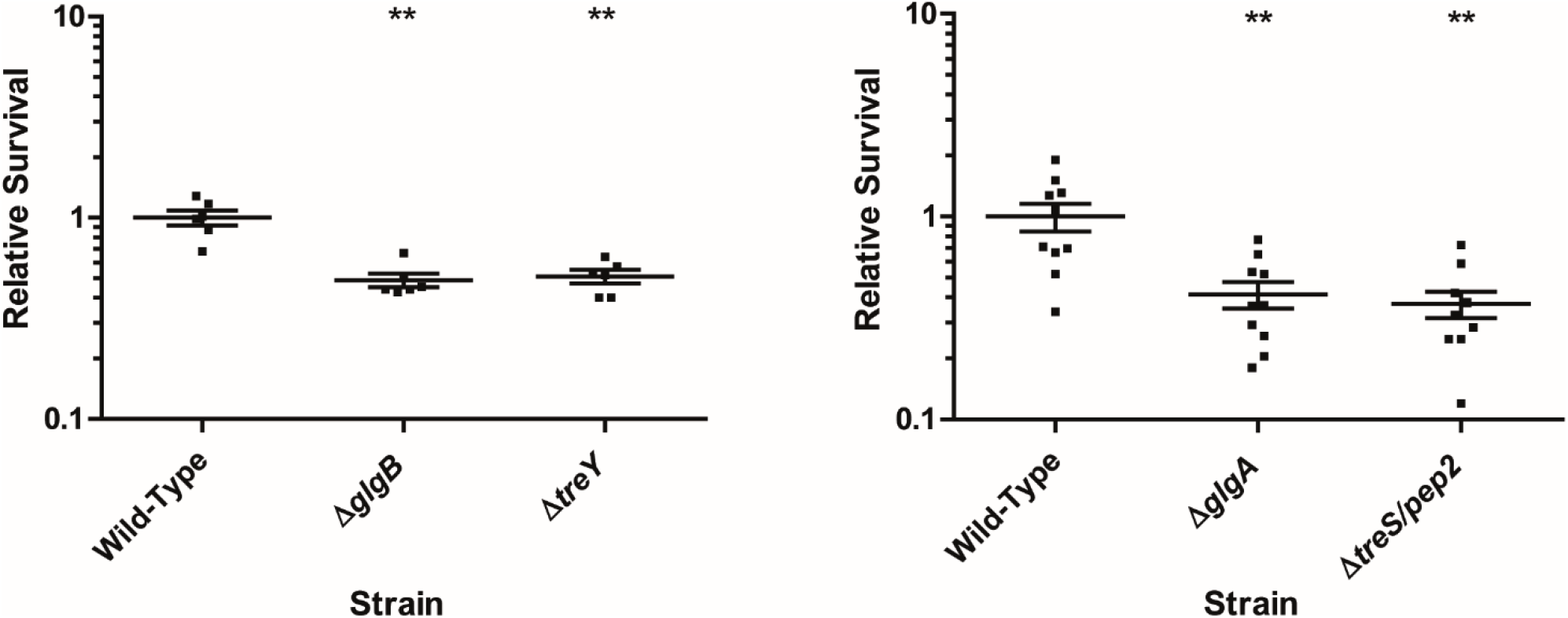
Survival of PA01 mutant strains on a steel work surface. Scatter graphs showing recovered CFUs for PA01 mutants after 24h incubation on a sterile brushed steel work surface, relative to WT PA01. X-axis shows different *tre*/*glg* mutants, Y-axis shows recovered CFUs as a fraction of the mean WT value. Individual data points are shown, with mean values ± standard error indicated with bars. Significant differences from the WT are shown with asterisks (Mann-Whitney U test, **: p<0.01). Experiments were conducted at least twice independently for each mutant and representative assays versus WT PA01 are shown in each case.

## Discussion

The mechanisms that pathogenic bacteria use to tolerate external stresses are crucial to effective survival in the environment. In this study we characterise a central stress tolerance pathway in a major opportunistic pathogen. Using a combination of reverse genetics and analytical biochemistry, we have deciphered the configuration of the *P. aeruginosa* trehalose and α-glucan biosynthetic pathways and differentiated between the contributions these ubiquitous molecules make to desiccation and osmotic stress tolerance (Figure **1B**). To our knowledge, this is the first report characterising the mechanisms of trehalose and α-glucan biosynthesis and their physiological functions in an organism possessing both the GlgA and GlgE pathways.

GlgA catalyses the first committed step in this pathway, polymerising UDP-glucose into linear α-glucan. This GlgA-α-glucan is then rapidly degraded into trehalose by TreY and TreZ. TreY catalyses the conversion of the reducing-end α-1,4 bond of α-glucan into an α-1, α-1 glycosidic bond, forming maltooligosyltrehalose. This molecule is then hydrolysed by TreZ, liberating trehalose. Unexpectedly, M1P was detected in strains lacking functional TreY and TreZ despite the loss of trehalose biosynthesis, suggesting an alternative route exists to biosynthesise maltose. Consequently, we established that MalQ disproportionates α-glucan with glucose to produce maltose. This result reconciles the surprisingly low activity of the TreS domain of TreS/Pep2 because it offers an alternative and more direct route from GlgA-derived α-glucan to maltose. The clustering of the genes for all four enzymes in this part of the pathway would allow for coordinated transcriptional regulation. Since the production of maltose by MalQ is dependent on the presence of glucose, the configuration of the pathways in PAO1 with the kinetic bottleneck in TreS offers a potential advantage. When glucose is limiting, it would be expected that trehalose would be the main metabolite accumulated because neither MalQ nor TreS would be capable of generating much maltose. However, when glucose is abundant, much of the flux towards trehalose would be diverted to maltose via MalQ. The consequence is that GlgE-derived branched α-glucan could then accumulate.

Once M1P has been produced by TreS/Pep2 and polymerised into linear α-glucan by GlgE, GlgB introduces α-1,6 branch points to the linear chain to form branched α-glucan. Iterative rounds of extension and branching leads to the final product of the GlgE pathway. The genes for the three enzymes of this part of the pathway are clustered. Branched α-glucan is recycled by the phosphorylase GlgP, which degrades the non-reducing ends of α-glucan chains producing G1P, and the debranching enzyme GlgX. Due to steric constrains in the vicinity of branch points, phosphorylases such as GlgP are unable to degrade external chains with a DP of 4 or less [44]. However, we have shown that GlgX hydrolyses α-1,6 linked external branches with a preference for DP 4. It is therefore possible to generate branched polymer in the presence of GlgX, the gene for which is clustered with *glgA*, *treY*, *treZ* and *malQ*, provided the external chains have a DP > 4. Thus, PAO1 is capable of fully degrading branched α-glucan into glucose-1-phosphate when the orphaned gene *glgP* is expressed, allowing GlgP and GlgX to work together.

The GlgA and GlgE proteins in PA01 and other *Pseudomonas* spp. enable two alternative mechanisms for linear α-glucan biosynthesis. We were unable to detect structural or functional differences between the α-glucan chains produced by either enzyme, with both molecules amenable to branching by GlgB and able to confer desiccation tolerance. On first inspection, the reasons for this apparent redundancy are unclear. However, despite producing α-glucan as an intermediate molecule, the cluster of genes including *glgA* is entirely geared towards the production of either trehalose or maltose *in vivo*. TreS/Pep2 deletion leads to an accumulation of intracellular trehalose, but this is not matched by a corresponding increase in GlgA-derived α-glucan due to rapid flux through TreY/TreZ that is assisted by a favourable equilibrium towards hydrolysis in the last step. This suggests that the vast majority of the α-glucan accumulated by *P. aeruginosa* will be produced by GlgE.

Trehalose and α-glucan biosyntheses in *Pseudomonas* spp. are clearly more closely integrated than previously suspected. Our analysis overturns earlier models for trehalose synthesis in *Pseudomonas* based on discrete sources of glycogen and maltose, with the source of the latter being previously unclear. Instead trehalose appears to be an intermediate that is both produced from and integral to the production of α-glucan via maltose (Figure **1B**). Consequently, the intracellular levels of both molecules must be linked, with their relative abundance dependent both on the level of the other molecule and on the relative activity of the respective catabolic and anabolic enzymes. Controlling the balance between trehalose and α-glucan biosynthesis in *P. aeruginosa* is clearly central to an effective bacterial response to water stresses. In previous studies, trehalose has been extensively implicated in bacterial responses to both osmotic and desiccating conditions [47, 59–62]. Conversely, α-glucan has been very rarely associated with bacterial stress tolerance, with published roles in carbon and energy storage [63] and virulence [64]. Based on our revised understanding of the relationship between trehalose and α-glucan biosynthesis, we revisited the roles of both molecules in mediating water stress phenotypes.

All tested PA01 mutants that lacked the ability to synthesise trehalose were osmostress sensitive, confirming a role for trehalose in osmotic stress tolerance. This was irrespective of the α-glucan concentration, because the α-glucan over-accumulating Δ*treY* and Δ*treZ* mutants were both sensitive. While this suggests that α-glucan may not play a direct role in tolerance to osmostress, a PAO1 Δ*glgB* mutant lacking the α-glucan branching enzyme was also osmostress sensitive, despite its trehalose level exceeding those of wild-type PA01 under unstressed conditions. It is possible that α-glucan may function as a substrate to ensure optimal trehalose production in response to osmostress. Alternatively, insoluble glucan functions as a major carbon sink. This may have wider metabolic and physical consequences for the cell, including osmostress sensitivity.

Although trehalose and α-glucan are biosynthetically linked, they have distinct roles in the protection of *Pseudomonas* spp. during water stress. By genetically decoupling cellular responses to distinct water stresses, we showed that trehalose protects *Pseudomonas* spp. against osmotic stress, most likely due to its role as a compatible solute [18]. However, and contrary to previous reports, trehalose does not directly protect cells against desiccation. This function is instead attributed to the polysaccharide α-glucan. While the presence of trehalose correlates with protection against desiccation stress, this is only true when the cell retains the ability to process trehalose into α-glucan *via* the GlgE pathway. The loss of α-glucan biosynthesis resulted in desiccation sensitive strains, irrespective of the ability to synthesise trehalose. When the levels of intracellular GlgA- and GlgE-derived linear α-glucans are increased in the Δ*glgB* strain desiccation resistance was also increased. This suggests that α-glucan protects PAO1 against desiccation stress irrespective of branching. Intriguingly, despite the distinct stress response roles played by trehalose and α-glucan in *P. aeruginosa*, disruption of either biosynthetic pathway significantly compromised abiotic surface survival. This suggests that *Pseudomonas* survival in the environment relies on the integrated deployment of both stress tolerance molecules. This presents an attractive rationale for the close genetic integration of trehalose and α-glucan biosynthesis in *P. aeruginosa* and raises the possibility of targeting the GlgA/GlgE pathways to limit pathogen survival in clinical and other relevant settings.

Our findings have implications for models of trehalose metabolism and function in many bacterial species. GlgE pathway genes are present in around 14 % of all sequenced bacterial genomes, including *Pseudomonas* spp. [45, 65, 66]. In the first instance, cell behaviours previously attributed to trehalose could in fact be mediated by α-glucan. For example, the desiccation tolerance phenotype attributed to trehalose in *Bradyrhizobium japonicum* [23] may in fact be an α-glucan phenotype, as the *B. japonicum* genome contains a complete *glgE* operon. Similarly, trehalose has been implicated as a virulence factor in *Pto* [47] and PA14 [48]. However, the mutations in both cases would also lead to abolition of α-glucan biosynthesis, raising the possibility that α-glucan plays a role in *Pseudomonas* virulence in a similar manner to *M. tuberculosis* [25].

There are still several outstanding questions concerning the role of trehalose and α-glucan in *P. aeruginosa*. At this stage the cellular localisation of α-glucan in PAO1 is unclear. *Pseudomonas fluorescens* has been shown to produce extracellular α-glucan [67] and mycobacterial α-glucan localises to both the cytosol and the extracellular capsule [34]. It is feasible that α-glucan acts as an additional exopolysaccharide that contributes to PA01 desiccation stress tolerance, although this remains to be investigated. The regulation of trehalose and α-glucan synthesis and deployment largely remain to be determined, as does the integration of their deployment with other stress response pathways and compatible solutes. The GlgA and GlgE operons are upregulated in both PAO1 and *P. syringae Pto* under conditions associated with osmotic stress [62, 68], and during water stress in *P. putida* KT2440 [61], although the signalling pathways responsible for this are currently undefined.

There is the potential for a futile cycle involving linear α-glucan being converted to trehalose and/or maltose and subsequently repolymerised to α-glucan at the expense of ATP. The main mechanism preventing this happening with branched polymer would be the requirement for GlgP activity to allow GlgX to debranch the polymer. However, linear α-glucan could also be cycled. The potential mechanisms to avoid this include a separation of GlgA and GlgE-derived linear α-glucan by time, space, or DP. It is feasible, for example, that short maltooligosaccharides are preferentially degraded by TreY and TreZ and that longer ones are preferentially branched by GlgB (e.g. DP 14 [55]). This might be kinetically controlled by GlgA acting slowly and GlgE acting quickly and/or substrate specificity. In addition, it is possible that GlgE can extend branched polymers but GlgA cannot. Even with equal enzyme activities, GlgE would elongate at twice the rate, two glucose residues at a time. Note that we have shown that GlgE elongates both odd and even acceptor substrates, so its products would not have an exclusively even length. These issues require additional research.

Finally, the importance of trehalose and α-glucan during human infection by PAO1 is unclear. Although not completely understood, capsular α-glucan is known to be essential for the full virulence of *M. tuberculosis* in a murine model [35]. This involves the binding of α-glucan to phagocytic cells via complement receptor 3, the blocking of dendritic cell function, and interactions with the C-type lectin DC-SIGN. PA14 *tre-* mutants were not attenuated in growth in a range of animal infection models [48]. However, these models are unlikely to reflect the significant osmolarity and desiccation stresses present in chronically infected CF lungs, or in the environmental reservoirs that contribute to the prevalence of community and hospital-acquired infections.

## Materials and Methods

### Strains and growth media

Bacterial strains and plasmids used in this study are listed in Table **S1**. *Pseudomonas aeruginosa* PAO1 and *Escherichia coli* DH5α strains were routinely cultured in lysogeny broth (LB, [69]) at 37 °C, solidified with 1.5% w/v agar where appropriate. Tetracycline was included in plates/media at 12.5 μg/ml for *E. coli* and 100 μg/ml for *P. aeruginosa*. Gentamycin was used at 25 μg/ml. For NMR spectroscopy and stress tolerance assays, PA01 strains were grown in M9 medium (20 mM NH_4_Cl, 12 mM Na_2_HPO_4_, 22 mM KH_2_PO_4_, 8.6 mM NaCl, 1 mM MgSO_4_, 1 mM CaCl_2_, supplemented with 0.4% glucose, 0.4% casamino acids and 50 μM FeCl_3_) [69].

### Molecular microbiology techniques

Cloning was carried out using standard molecular biology techniques [70]. Polymerase chain reaction (PCR) products and plasmid digestion derivatives were purified using NucleoSpin^®^ Gel and PCR Clean-up kits (Macherey-Nagel). Plasmids were extracted from *E. coli* DH5α strains using the NucleoSpin^®^ Plasmid DNA Purification kit (Macherey-Nagel). Cloning was performed using the appropriate restriction endonucleases and T4 DNA ligase (New England Biolabs). PCR reactions were conducted using Q5^®^ High-Fidelity DNA polymerase (New England Biolabs). Primary denaturation was performed for 30 s at 98 °C. Following this, 30 cycles of denaturation (98 °C, 10 s), annealing (varying temperatures, 30 s) and extension (72 °C, 30 s) was performed, followed by a final extension at 72 °C for 120 s. The annealing temperature for a given reaction was calculated as Tmb – 2 °C, where Tmb represents the lowest Tm of the primer pair. The sequences and melting temperatures (Tm) of primers used in this study are listed in Table **S2**.

### Genetic manipulation of PA01

Gene deletion vectors were constructed by amplifying the upstream and downstream regions flanking the desired gene from the PAO1 genome using primers 1-54 listed in Table **S2**. Amplified up- and downstream flanking regions were then cloned into the multiple cloning site of the suicide vector pTS1 [71]. PA01 cells were transformed with the appropriate deletion vectors following the method described in [72]. Single crossover integrations of each plasmid into the chromosome were selected on tetracycline plates and purified by re-streaking onto fresh plates. Single colonies were picked and grown overnight in 200 ml LB medium without selection. Colonies where double crossovers had occurred were then selected by plating onto LB agar with 10 % w/v sucrose. Gene deletions were confirmed for tetracycline-sensitive, sucrose-resistant colonies by PCR using GoTaq^®^ (Promega) and primers listed in Table **S2**. The pME3087-derivative vector pΔ*PA2152* lacks the *sacB* counter-selection gene. In this case, we used tetracycline enrichment to select for double crossover candidates following the method described in [73] followed by PCR confirmation of successful gene deletions as described above.

To introduce *otsA/otsB* genes into the PA01 *att::Tn7* site, the *otsA/otsB* locus was amplified from the *Microbacterium koreensis* 3J1 genome [58] using primers 55 and 56. The resulting PCR product was cloned into the multiple cloning site of pME6032 [74]. Primers 57 and 58 were then used to amplify *otsA/otsB* in addition to the *tac* promoter and terminator of pME6032. This product was then cloned into the pUC18-mini-*Tn7*T-Gm vector [75] and used to transform PA01 via electroporation. To overexpress *glgB in trans*, the *glgB* open reading frame was amplified with primers 69 and 70 and cloned into plasmid pME6032 [74] between the *Eco*RI and *Xho*I sites. pME-*glgB* was then transferred into PA01 strains by electroporation.

### Enzymology

Unless otherwise stated, the genes coding for relevant enzymes were amplified from PA01 genomic DNA by PCR with primers 58-68 & 71-72 and sub-cloned into a pET101 Directional TOPO vector (Invitrogen) allowing protein production with an N-terminal 6×His-tag. *E. coli* SoluBL21 cells (AMS Biotechnology Europe Ltd) bearing each resultant plasmid were grown in Lysogeny Broth to an OD_600_ of 0.6 (unless otherwise state) with agitation and recombinant gene expression was induced with 0.5 mM isopropyl β-D-thiogalactopyranoside (IPTG). Recombinant protein was purified using a 1 ml HisTrap FF column (GE Healthcare, Amersham, United Kingdom) with imidazole gradient elution. Purified proteins were buffer exchanged into 100 mM bis-tris propane, pH 7.0, containing 50 mM NaCl, unless otherwise stated.

The expression of recombinant PAO1 *glgA* was induced with IPTG and 0.4 % dimethyl sulfoxide (DMSO). The culture was incubated for a further 5 hours at 37 °C with agitation. Purified GlgA was incubated with 10 mM NDP-glucose for 2 days at 20 °C. The product mixture was analysed using MALDI-TOF mass spectrometry by adding 1:1 (v/v) matrix (20 mg/ml di-hydroxy-benzoic acid in 50:50 MeCN/H_2_O) before spotting onto a target plate [33]. ^1^H-NMR spectra were recorded as described below using samples that contained 10% (v/v) D_2_O.

Recombinant PAO1 *glgP* was induced with IPTG and the culture was grown for a further 16 hours at 18 °C. Purified GlgP was buffer exchanged into 20 mM Hepes, pH 7.5. GlgP was incubated with 10 mM maltohexaose and 1 mM pyridoxal phosphate in 50 mM phosphate buffer, pH 7.0 at 20 °C for 18.5 hours. The reaction mixture was analysed using NMR spectroscopy.

Rather than cloning the PAO1 *treS/pep2* gene from genomic DNA, it was synthesised (Genscript Corporation, Piscataway, NJ, U.S.A.) with optimum codon usage for expression in *E. coli*, allowing the production of the enzyme with an N-terminal 6×His-tag with a TEV cleavage site. The synthetic gene was cloned into a pET-21a(+) (Novagen, Watford, United Kingdom) vector using *Nde*I and *BamH*I restriction sites. The expression of the recombinant enzyme in *E. coli* BL21(DE3) was induced with IPTG and 0.4 % dimethyl sulfoxide (DMSO). The culture was incubated for a further 18 h at 18 °C with agitation. The protein was purified using a nickel-affinity column followed by an S200 16/60 gel filtration column (Pharmacia Biotech, Amersham, United Kingdom) eluted with 20 mM HEPES buffer, pH 7.5, containing 10% v/v glycerol and 200 mM NaCl. The maltokinase activity of TreS-Pep2 was monitored using a continuous enzyme-coupled spectrophotometric assay following the release of ADP. Chemicals were purchased from Sigma Aldrich. Unless otherwise stated, all enzyme assays were carried out at 21 °C in 50 mM bis-tris propane, pH 8.0, containing 5 mM MgCl_2_, 0.3 mM NADH, 1 mM phosphoenolpyruvate, 1 U lactate dehydrogenase, 1 U pyruvate kinase and 0.2 mg/ml bovine serum albumin. Saturation kinetics for ATP and maltose were measured in a 6 × 8 matrix in a Nunc 384 well plate using a BMG Clariostar plate reader. Six maltose concentrations were used from 0.125 to 4.0 mM and eight ATP concentrations from 8 μM to 1.0 mM. The effect of pH was measured using 50 mM buffers; bis-tris (pH 6.0) or bis-tris propane (pH 7.0, 8.0 and 9.0) with 1 mM ATP and 2 mM maltose. The temperature-dependence of activity between 25 and 50 °C was determined using 1 mM ATP and 4 mM maltose using Perkin Elmer Lambda 25 or 35 spectrophotometers fitted with Peltier temperature controllers. Enzyme concentrations were such to allow reactions to progress linearly for 5 min (cuvette) or 100 s (plate reader) with total donor consumption being < 10%. Initial rates (v_0_/[E]) were determined by monitoring the oxidation of NADH at 340 nm.

Kinetics curves were fitted to a random order bi-substrate ternary complex model (Equation 1) using global fitting in GraphPad Prism 8. The value for *K*_*mA*_ was obtained by repeating the fitting with the all of the terms for A and B reversed, given the symmetry of the mechanism.

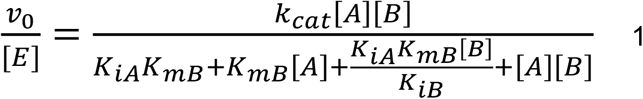

Recombinant PAO1 *glgE* was induced with IPTG and the culture was grown for a further 5 hours at 37 °C. GlgE activity was measured using a spectrophotometric stopped assay that detected the release of inorganic phosphate using Malachite green [33]. The effect of pH was measured using 20 mM buffers; bis-tris (pH 6.0) or bis-tris propane (pH 7.0, 8.0 and 9.0) with 0.25 mM M1P and 1 mM maltohexaose. The effect of temperature was measured with 1 mM each of M1P and maltohexaose. The default GlgE assays were carried out at 30 °C and a 1 mM fixed substrate concentration. Disproportion activity was detected using MALDI mass spectrometry.

Individual kinetic curves were fitted to either a Michaelis-Menten model (Equation 2) or one modified to include substrate inhibition (Equation 3) using GraphPad Prism 8.

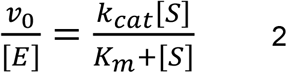

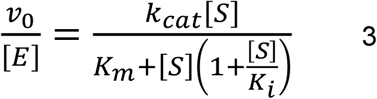

Recombinant PAO1 *malQ* was induced with IPTG and the culture was grown for a further 16 hours at 18 °C. Purified MalQ was incubated with 1 mM substrate to test for disproportionation activity. Reaction mixtures were analysed using MALDI mass spectrometry. The production of maltose from maltohexaose was also tested using an assay coupled to the production of M1P and ADP by TreS/Pep2 as described above. Specifically, MalQ was incubated with TreS/Pep2, 1 mM ATP, 5 mM maltohexaose, 1 U lactate dehydrogenase, 1 U pyruvate kinase, 1 mM phosphoenol pyruvate, 0.3 mM NADH, 50 mM bis-tris propane, pH 7.0, and 5 mM MgCl_2_. Similarly, MalQ was incubated with TreS/Pep2, 1 mM maltohexaose, 4 mM ATP and 0.2 mg/ml bovine serum albumin, and the reaction mixture was analysed using NMR spectroscopy to detect the production of M1P.

Recombinant PAO1 *glgB* was induced in auto-induction medium [76] and incubated for 5 hours at 37 °C. Purified GlgB was incubated with PAO1 GlgE, 1 mM maltohexaose, and 10 mM M1P for 2 days at 20 °C. The resulting α-glucan particles were analysed using transmission electron microscopy after staining with 2% aqueous uranyl acetate, pH 4.5 [55]. Samples were also buffer exchanged into sodium acetate buffer, pH 4.0, using 10 kDa cut-off filters that removed maltohexaose and M1P. The washed particles were digested with 2 U of isoamylase for 3 hours at 37 °C [55]. The resulting product mixtures were analysed using MALDI mass spectrometry.

Recombinant PAO1 *glgX* was induced with IPTG and the culture was grown for a further 16 hours at 18 °C. Purified GlgX was incubated with 10 mg/ml rabbit liver glycogen in 20 mM bis-tris propane, pH 7.0. The resultant product mixture was analysed using MALDI mass spectrometry. Control reactions without GlgX contained no detectable maltooligosaccharides.

### Metabolite extraction

Overnight PA01 cultures were grown in M9 medium at 37 °C with shaking, then diluted to an OD_600_ of 0.5 in PBS. Mixed cellulose ester filter discs (Merck Millipore) were placed on the surface of M9 agar plates and coated with the diluted cell suspensions. Plates were then incubated at 37 °C for 16 hours before cells were harvested from each disc and resuspended in 5 ml of ddH_2_O in a 15 ml plastic tube by vigourous vortexing. Water was then removed from each sample by drying under vacuum for 16 hours. Following this step, sample dry weight was determined. Dried cells were resuspended in ddH_2_O and boiled at 95 °C for 20 minutes. Boiled cells were then centrifuged (10,000 × g, 10 minutes) and the resulting supernatant was subject to additonal boiling and centrifugation steps. The supernatant, now containing the total soluble metabolite content of the cell sample, was dried under vacuum for a further 16 hours. The resulting dried pellet was resuspended in 1200 μl deuterium oxide (D_2_O) and analysed by NMR spectroscopy.

### Nuclear magnetic resonance spectroscopy

Metabolite samples were subjected to ^1^H-NMR spectroscopy using a Bruker AVANCE III 400 spectrometer. Samples were examined at room temperature using 400 MHz with water supression. Metabolite chemical shifts were recorded as parts per million (ppm) compared to the standard: 0.5 mM trimethylsilyl propanoic acid (TMSP; 0.00 ppm). Spectra were analysed using Topspin 3.0 (Bruker) and the concentrations of metabolites were established through the manual integration of peaks relative to TMSP. The resonances were assigned based on previously established spectra [77].

### α-Glucan staining

The production of α-glucan in PA01 colonies was visualised with iodine crystals (Sigma-Aldrich). A few crystals were placed in the lids of inverted plates upon which bacterial strains were streaked. Strains were placed in a fume hood and exposed to sublimating iodine vapour for 2 minutes and were immediately photographed. α-Glucan in metabolite extracts was visualised by adding 15% (v/v) Lugol’s solution (Sigma-Aldrich).

### Osmotic stress assays

The growth kinetics of each bacterial strain were measured using a 96-well microplate spectrometer (Biotek Instruments). Overnight PA01 cultures were grown in M9 medium (20 mM NH_4_Cl, 12 mM Na_2_HPO_4_, 22 mM KH_2_PO_4_, 8.6 mM NaCl, 1 mM MgSO_4_, 1 mM CaCl_2_, and 0.4% glucose, 0.4% casamino acids, 50 μM FeCl_3_) at 37 °C with shaking, then diluted to OD_600_ of 0.01 in M9 medium. Five μl of each diluted sample was used to inoculate 150 μl of M9 medium. To examine osmotic stress conditions, growth media were supplemented with 0.85 M NaCl. Growth was monitored by measuring the OD_600_ every hour until stationary phase had been attained. Assays were conducted in triplicate and repeated at least twice independently, with a representative sample shown in each case.

### Desiccation tolerance assays

Overnight PA01 cultures were grown in M9 medium at 37 °C with shaking, then diluted to an OD_600_ of 0.1 in PBS. Ten μl of each culture were spotted onto 15 mm grade 1 Whatman filter discs (GE Healthcare Life Sciences). After drying for 1 minute at room temperature, discs were placed onto M9 agar plates and incubated at 37 °C for 4 hours to enable bacteria to recover and begin dividing. After incubation, the filter discs were subjected to controlled desiccation in tightly sealed bell chambers containing either water (100% RH) or a saturated solution of NaCl (75% RH [78]) for 2 hours. Bacteria were then recovered from filter discs in 3 ml of PBS and serially diluted onto LB agar plates. The number of colony-forming units (CFU) were determined for each strain. Assays were conducted in triplicate (technical replicates) and repeated at least twice independently on different dates (biological replicates).

### Surface survival assays

Overnight PA01 cultures were grown in LB medium at 37 °C with shaking, washed once with PBS, then diluted to an OD_600_ of 1.0 in PBS. The number of colony-forming units (CFU) was determined for each strain by serial dilution onto LB agar plates. Ten μl of each culture were spotted onto a sterile, brushed steel work surface and allowed to dry. Spots were incubated for 24 h at room temperature and in ambient atmospheric conditions (Norwich, UK, August 2020). Bacteria were then recovered in 10 μl of PBS and serially diluted onto LB agar plates. The number of colony-forming units (CFU) was determined for each strain and normalised to the CFU of the initial inoculum. Assays were conducted with at least six replicates per sample and repeated twice independently.

### Statistical analysis of desiccation data

Log_10_(CFU) values from the desiccation assays were analysed by linear mixed modelling using restricted maximum likelihood. The random effect was Date of experiment and the fixed effects were Condition*Strain, where * is the crossing operator. Condition included the initial culture (T_0_) and growth at 100% RH and 75% RH. The three technical replicate samples with the same Condition and Strain tested on the same Date provided the residual error term. The statistical significance of Condition, Strain and their interaction was determined by F-tests of Wald statistics. The response of a strain to lower humidity was calculated as the difference between its predicted mean log_10_(CFU) at 100% RH and at 75% RH. The standard error of the difference (SED) between the responses of mutant and wild-type strains was calculated from the appropriate standard errors (SE) with the formula: SED = (SE[100%, Mutant]^2^ + SE[75%, Mutant]^2^ + SE[100%, wild-type]^2^ + SE[75%, wild-type]^2^)^0.5^. The significance of the response difference was tested using the normal distribution with a variance of SED^2^. Statistical significance between the desiccation responses of all PAO1 strains is summarised in Table **S3**.

---

**Figure S1:**
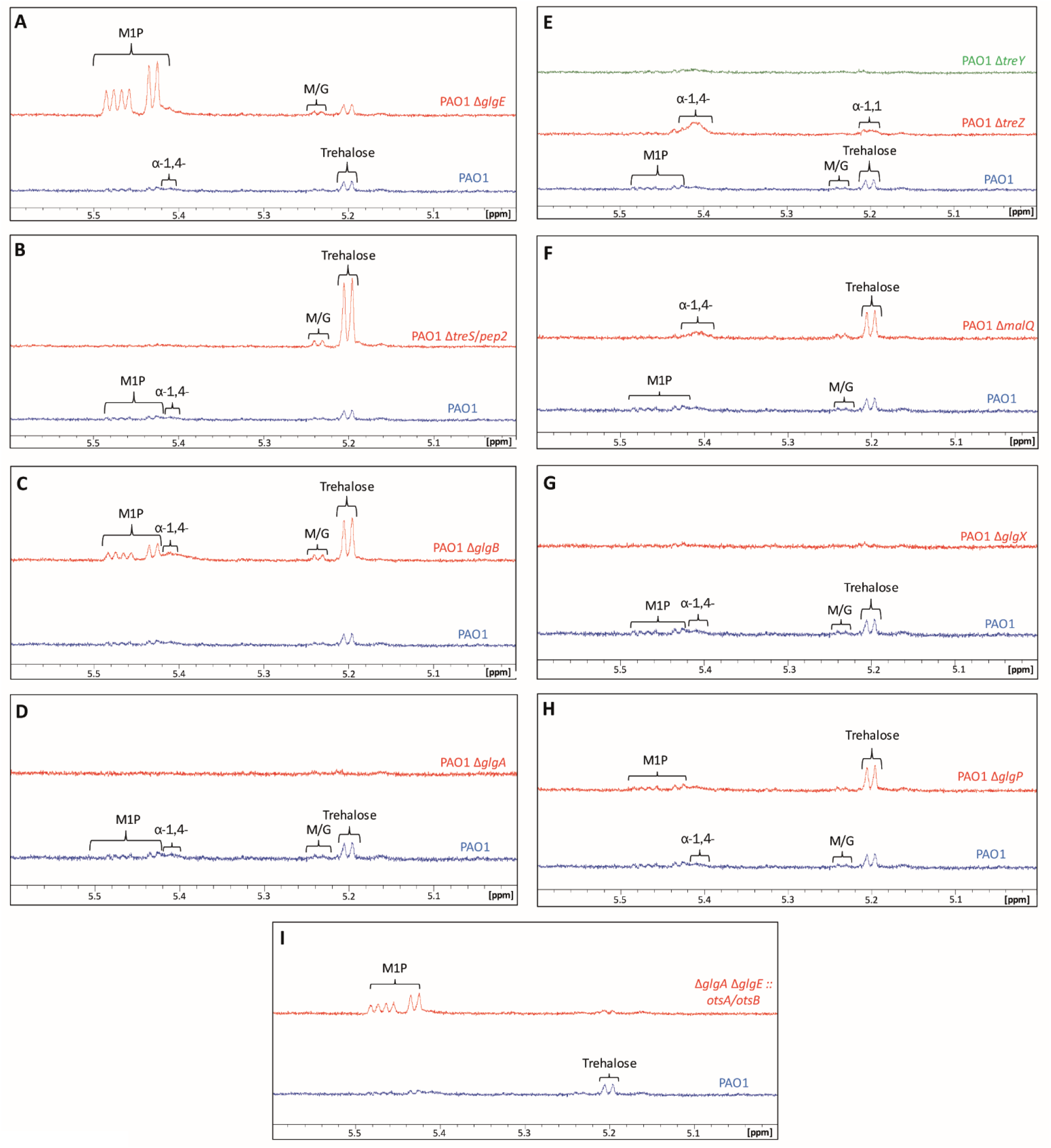
^1^H-NMR spectra for PA01*glgA* and *glgE* operon mutants. Peaks corresponding to key metabolites: M/G – maltose/glucose, α-1,4-– α-glucan, α-1,1-– probable terminal linkage of maltooligosyltrehalose. Mutant strains are labelled as follows: **A)** Δ*glgE*, **B)** Δ*treS/pep2*, **C)** Δ*glgB*, **D)** Δ*glgA*, **E)** Δ*treZ*/Δ*treY*, **F)** Δ*malQ*, **G)** Δ*glgX*, **H)** Δ*glgP*, **I)** Δ*glgA* Δ*glgE :: otsA/otsB.*

**Figure S2:**
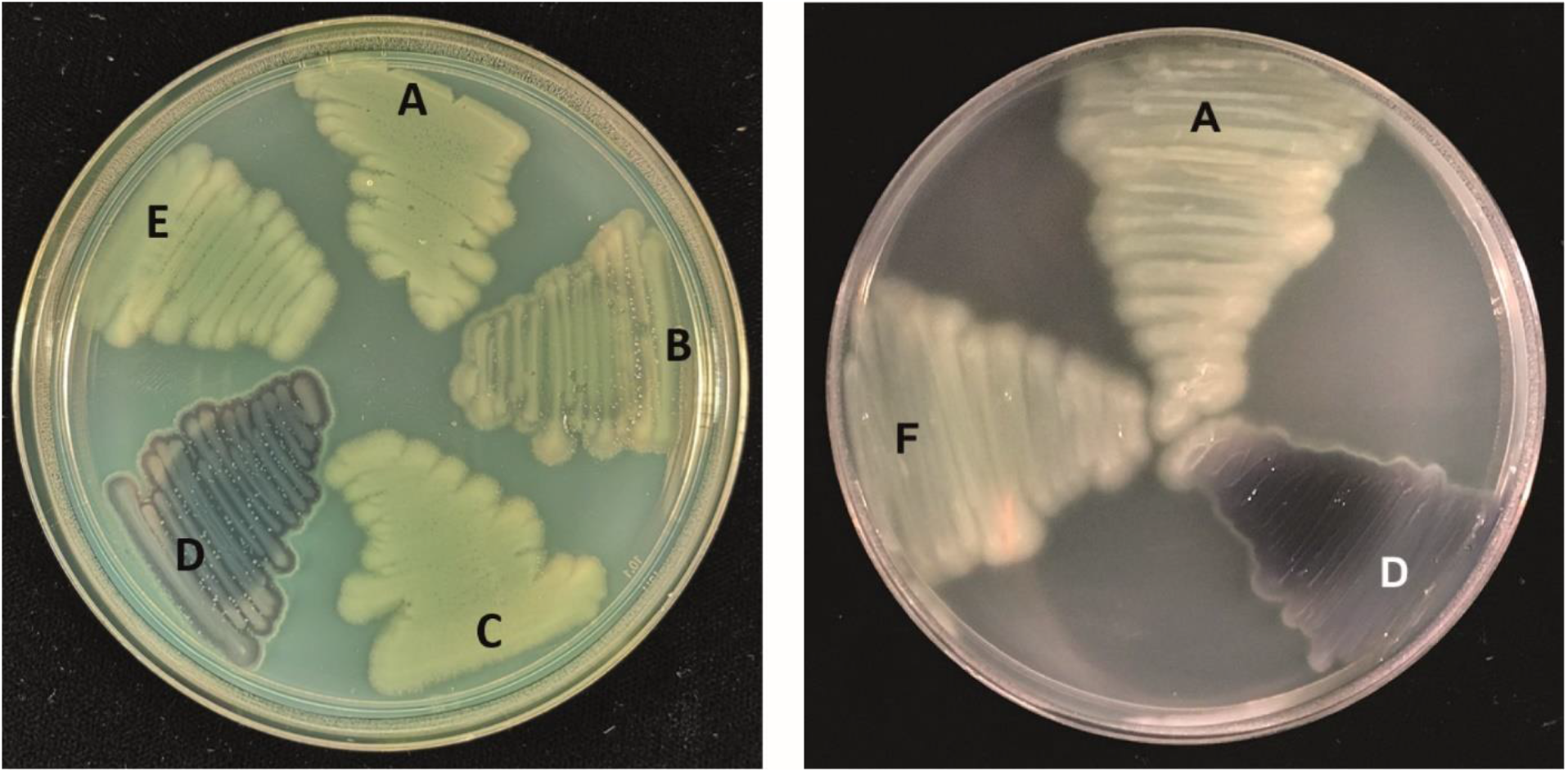
PA01 strains exposed to iodine vapour. Strains producing insoluble linear α-glucan stain blue-purple. Mutant strains are labelled as follows: **A)** Wild-type PAO1, **B)** Δ*glgB*, **C)** Δ*PA2151-53*, **D)** Δ*treY* Δ*PA2151-53* (PA01 Δ*4*), **E)** Δ*malQ* Δ*PA2151-53*, **F)** PA01 Δ*4* pME-*glgB*.

## Acknowledgements

We thank Sibyl Batey for assistance with data analysis.

---

**Table S1:** Strains and plasmids used in this study.

| Strain | Details | Source |
| --- | --- | --- |
| ***P. aeruginosa*** |  |  |
| PAO1 | Wild-type *Pseudomonas aeruginosa* PAO1 | [79] |
| PAO1 $\Delta glgE$ | Non-polar deletion of *PA2151* | This study |
| PAO1 $\Delta treS/pep2$ | Non-polar deletion of *PA2152* | This study |
| PAO1 $\Delta glgB$ | Non-polar deletion of *PA2153* | This study |
| PAO1 $\Delta glgA$ | Non-polar deletion of *PA2165* | This study |
| PAO1 $\Delta treZ$ | Non-polar deletion of *PA2164* | This study |
| PAO1 $\Delta malQ$ | Non-polar deletion of *PA2163* | This study |
| PAO1 $\Delta treY$ | Non-polar deletion of *PA2162* | This study |
| PAO1 $\Delta glgX$ | Non-polar deletion of *PA2160* | This study |
| PAO1 $\Delta glgP$ | Non-polar deletion of *PA2144* | This study |
| PAO1 $\Delta glgA/E$ | Non-polar deletion of *PA2165* & *PA2151* | This study |
| PAO1 $\Delta 4$ | Non-polar deletion of *PA2162* & *PA2151-PA2153* | This study |
| PAO1 :: *otsA/otsB* | PAO1 containing *p<sub>tac</sub>-otsA/B* at the *att::Tn7* insertion site, Gm<sup>R</sup> | This study |
| PAO1 $\Delta treS/pep2$ :: *otsA/otsB* | PAO1 $\Delta treS/pep2$ containing *p<sub>tac</sub>-otsA/B* at the *att::Tn7* insertion site, Gm<sup>R</sup> | This study |
| PAO1 $\Delta glgA$ :: *otsA/otsB* | PAO1 $\Delta glgA$ containing *p<sub>tac</sub>-otsA/B* at the *att::Tn7* insertion site, Gm<sup>R</sup> | This study |
| PAO1 $\Delta glgA/E$ :: *otsA/otsB* | PAO1 $\Delta glgA/E$ containing *p<sub>tac</sub>-otsA/B* at the *att::Tn7* insertion site, Gm<sup>R</sup> | This study |
| ***E. coli*** |  |  |
| BL21-(DE3) pLysS | Sm<sup>R</sup>, K12 *recF143 lacI<sup>q</sup> lacZΔ.M15, xylA, pLysS* | Novagen |
| DH5α | *endA1, hsdR17(r<sub>K</sub>-m<sub>K</sub>+), supE44, recA1, gyrA (Nal<sup>r</sup>), relA1, $\Delta(lacIZYA-argF)$ U169, deoR, $\Phi 80dlac\Delta(lacZ)M15$* | [80] |
| **Plasmids** |  |  |
| pTS1 | Tet<sup>R</sup>, suicide vector; *ColE1*-replicon, *IncP-1*, *Mob*, *lacZ* | [81] |
| pTS1-*glg/tre* vectors | pTS1 with $\Delta glg/\Delta tre$ constructs as *Bam*HI-*Hind*III inserts | This study |
| TOPO101 | Amp<sup>R</sup>, pET101 directional TOPO vector, N-term His<sub>6</sub>-tag | Invitrogen |
| pET21a(+) | Amp<sup>R</sup>, purification vector, C-terminal His<sub>6</sub>-tag | Novagen |
| TOPO101-*glg/tre* vectors | TOPO101 with *glg/tre* genes inserted | This study |
| pET21a-*treS/pep2* | pET21a with *treS/pep2* gene as *Nde*I-*Bam*HI insert | This study |
| pME3087 | Tet<sup>R</sup>, suicide vector; *ColE1*-replicon, *IncP-1*, *Mob* | [82] |
| pME3087-*treS/pep2* | pME3087 with *treS/pep2* as an *EcoRI-BamHI* insert | This study |
| pME6032 | Tet<sup>R</sup>, P<sub>K</sub>, 9.8 kb pVS1 derived shuttle vector | [74] |
| pME-*glgB* | pME6032 with PA01 *glgB* as an *EcoRI-XhoI* insert | This study |
| pUC18T-mini-Tn7T-Gm | Amp<sup>R</sup>, Gm<sup>R</sup>, Tn7 insertion vector | [83] |
| pUC18-mini-Tn7TGm-*otsA/B* | pUC18-miniTn7Gm with *p<sub>tac</sub>-otsA/B* fragment as a *HindIII-SpeI* insert. | This study |
| pTNS2 | Helper plasmid for *att::Tn7* insertion | [83] |

**Table S2:**
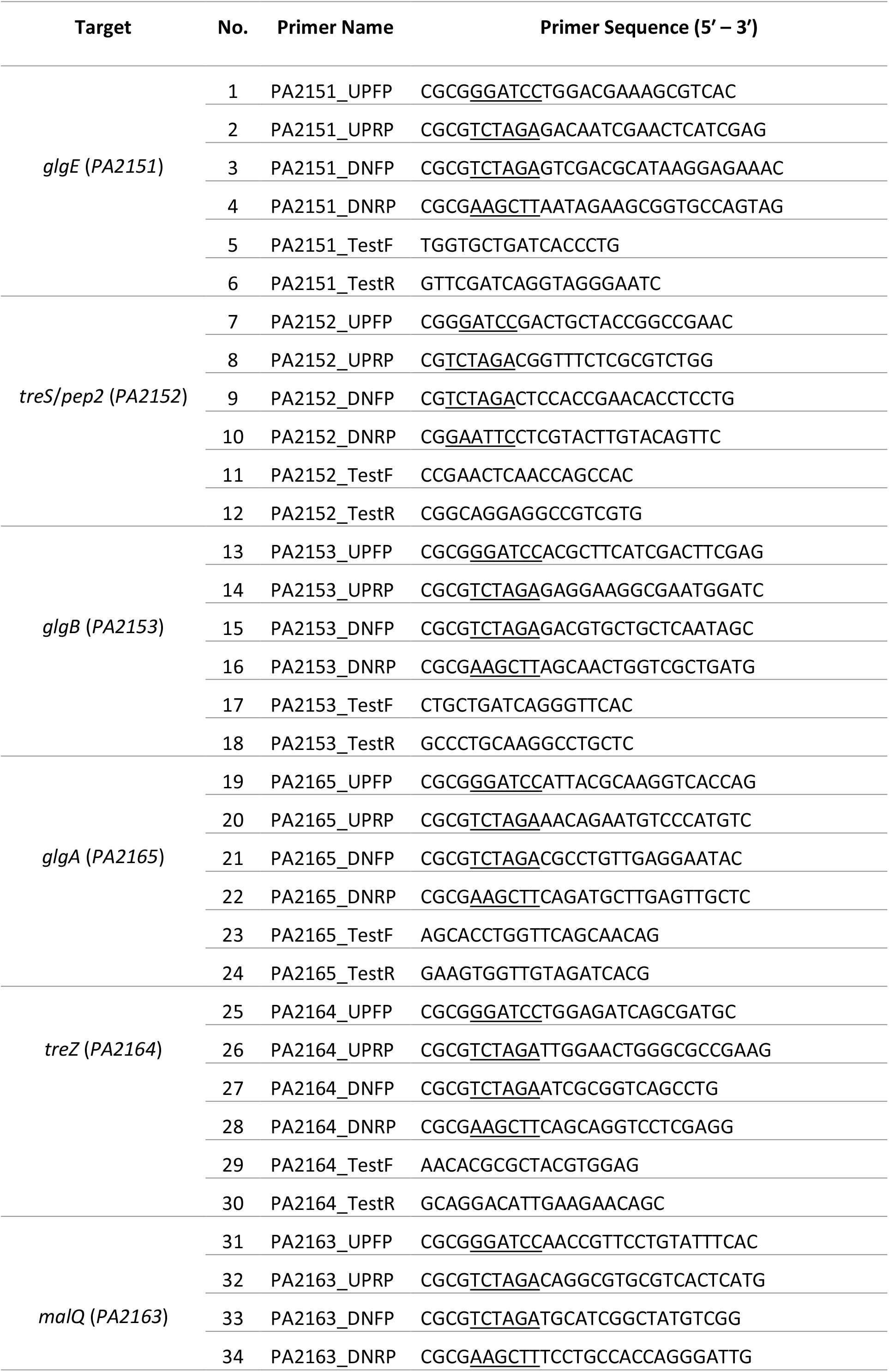
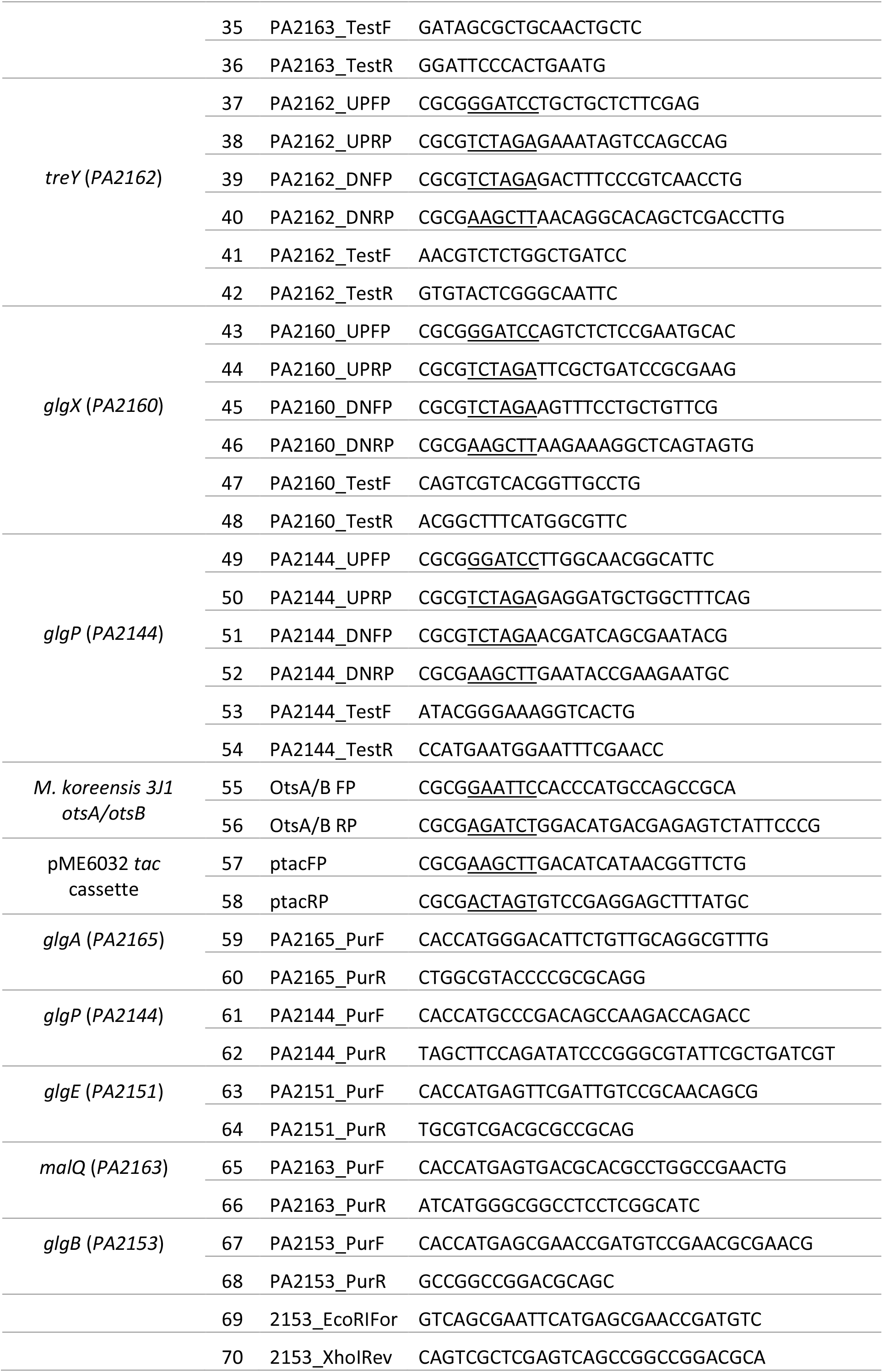
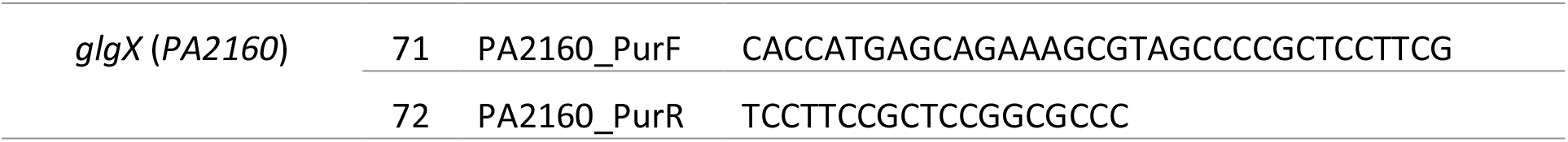
Primers used in this study.

**Table S3:** Statistical significance between the desiccation responses of PA01 and mutant strains. Predicted means and standard errors of log_10_(CFU/ml) for desiccation assays in Figures 9 and 10, calculated by linear mixed modelling (see text for details). The response to desiccation is the difference between log_10_(CFU) at 100% and 75% RH. P-values were calculated by Z-tests using the normal distribution.

| Strain | Condition |  |  | Difference |
| --- | --- | --- | --- | --- |
|  | 100% | 75% | T<sub>0</sub> | 100% – 75% |
| WT PA01 | 7.5820 | 6.4816 | 6.2734 | 1.1004 |
| PA01 :: otsA/B | 7.8715 | 5.6518 | 6.4538 | 2.2197 |
| $\Delta\text{alg}$ | 7.7245 | 4.7386 | 5.5958 | 2.9859 |
| $\Delta\text{glgA}$ | 7.3640 | 4.7875 | 6.2233 | 2.5765 |
| $\Delta\text{glgA} :: \text{otsA/B}$ | 7.7361 | 6.8782 | 6.4444 | 0.8579 |
| $\Delta\text{glgA} :: \text{otsA/B (+ 1 mM VA)}$ | 7.7312 | 3.8329 | 6.2089 | 3.8982 |
| $\Delta\text{glgA } \Delta\text{glgE} :: \text{otsA/B}$ | 7.5794 | 4.3929 | 6.1619 | 3.1866 |
| $\Delta\text{glgB}$ | 7.6090 | 7.3349 | 6.2340 | 0.2740 |
| $\Delta\text{treS/pep2}$ | 7.5332 | 4.3955 | 6.1655 | 3.1378 |
| $\Delta\text{treS/pep2} :: \text{otsA/B}$ | 7.6880 | 3.7468 | 6.3850 | 3.9412 |

| Strain | Condition |  |  | SE of response |
| --- | --- | --- | --- | --- |
|  | 100% | 75% | T<sub>0</sub> | 100% – 75% |
| WT PA01 | 0.0909 | 0.0909 | 0.0949 | 0.1286 |
| PA01 :: otsA/B | 0.2625 | 0.2625 | 0.2625 | 0.3713 |
| $\Delta\text{alg}$ | 0.3224 | 0.3224 | 0.3224 | 0.4559 |
| $\Delta\text{glgA}$ | 0.1859 | 0.1859 | 0.2276 | 0.2629 |
| $\Delta\text{glgA} :: \text{otsA/B}$ | 0.1608 | 0.1608 | 0.1608 | 0.2274 |
| $\Delta\text{glgA} :: \text{otsA/B (+ 1 mM VA)}$ | 0.2628 | 0.2628 | 0.2628 | 0.3716 |
| $\Delta\text{glgA } \Delta\text{glgE} :: \text{otsA/B}$ | 0.2663 | 0.2663 | 0.2663 | 0.3766 |
| $\Delta\text{glgB}$ | 0.2635 | 0.2635 | 0.2635 | 0.3727 |
| $\Delta\text{treS/pep2}$ | 0.1725 | 0.1725 | 0.1725 | 0.2439 |
| $\Delta\text{treS/pep2} :: \text{otsA/B}$ | 0.1859 | 0.1859 | 0.1859 | 0.2628 |

**Difference from WT PA01 in desiccation response**
| Strain | Difference of responses | SE of difference | P value |  |
| --- | --- | --- | --- | --- |
| PA01 :: otsA/B | 1.1194 | 0.3929 | 0.0044 | ** |
| $\Delta\text{alg}$ | 1.8855 | 0.4737 | <0.0001 | **** |
| $\Delta\text{glgA}$ | 1.4761 | 0.2926 | <0.0001 | **** |
| $\Delta\text{glgA} :: \text{otsA/B}$ | -0.2424 | 0.2612 | 0.35 | ns |
| $\Delta\text{glgA} :: \text{otsA/B (+ 1 mM VA)}$ | 2.7979 | 0.3932 | <0.0001 | **** |
| $\Delta\text{glgA } \Delta\text{glgE} :: \text{otsA/B}$ | 2.0862 | 0.3980 | <0.0001 | **** |
| $\Delta\text{glgB}$ | -0.8263 | 0.3942 | 0.036 | * |
| $\Delta\text{treS/pep2}$ | 2.0374 | 0.2758 | <0.0001 | **** |
| $\Delta\text{treS/pep2} :: \text{otsA/B}$ | 2.8408 | 0.2926 | <0.0001 | **** |

